# Comparing statistical and mechanistic models to identify the drivers of mortality within a rear-edge beech population

**DOI:** 10.1101/645747

**Authors:** Cathleen Petit-Cailleux, Hendrik Davi, François Lefèvre, Joseph Garrigue, Jean-André Magdalou, Christophe Hurson, Elodie Magnanou, Sylvie Oddou-Muratorio

## Abstract

Since several studies have been reporting an increase in the decline of forests, a major issue in ecology is to better understand and predict tree mortality. The interactions between the different factors and the physiological processes giving rise tree mortality, as well as the inter-individual variability in mortality risk, still need to be better assessed.

This study investigates mortality in a rear-edge population of European beech (*Fagus sylvatica* L.) using a combination of statistical and process-based modelling approaches. Based on a survey of 4323 adult beeches since 2002 within a natural reserve, we first used statistical models to quantify the effects of competition, tree growth, size, defoliation and fungi presence on mortality. Secondly, we used an ecophysiological process-based model (PBM) to separate out the different mechanisms giving rise to temporal and inter-individual variations in mortality by simulating depletion of carbon stocks, loss of hydraulic conductance and damage due to late frosts in response to climate.

The combination of all these simulated processes was associated with the temporal variations in the population mortality rate. The individual probability of mortality decreased with increasing mean growth, and increased with increasing crown defoliation, earliness of budburst, fungi presence and increasing competition, in the statistical model. Moreover, the interaction between tree size and defoliation was significant, indicating a stronger increase in mortality associated to defoliation in smaller than larger trees. Finally, the PBM predicted a higher conductance loss together with a higher level of carbon reserves for trees with earlier budburst, while the ability to defoliate the crown was found to limit the impact of hydraulic stress at the expense of the accumulation of carbon reserves.

We discuss the convergences and divergences obtained between statistical and process-based approaches and we highlight the importance of combining them to characterize the different processes underlying mortality, and the factors modulating individual vulnerability to mortality.

## Introduction

Global changes have been repeatedly reported to be the cause of forest decline and tree mortality, both in terms of background, non-catastrophic mortality (Van Mantgem et al. 2009, Lorenz and Becher 2012) and of massive, catastrophic mortality due to extreme, ‘pulse’ events (Allen et al. 2010; Lorenz and Becher 2012; Mueller et al. 2005). To predict how such a new regime of trees mortality will impact upon forest structure, composition and ecosystem services (Anderegg et al. 2015a; Choat et al. 2018), we need to better understand the respective roles of the various drivers and mechanisms underlying tree mortality.

Studying mortality poses several challenges, in particular because it is triggered by several factors and involves several interacting physiological processes. The factors triggering mortality include extreme, pulse climatic events (i.e. drought, storms, floods, heavy snow, late frosts, wildfires) or sudden changes in biotic interactions (i.e. emerging pests, invasive species), but also long-term climatic or biotic perturbations (i.e. recurrent water deficits, changes in competition at the community level) (Maraun et al. 2003; McDowell et al. 2011). Moreover, these factors can have interactive effects. For instance, drought may increase trees’ vulnerability to pests (Durand-Gillmann et al. 2014; Anderegg et al. 2015b) or predispose them to wildfires (Brando et al. 2014). Finally, a single factor triggering mortality may involve several underlying physiological processes, with several thresholds leading to mortality and potential feedback between them (McDowell et al. 2011). This is exemplified by drought, which is usually considered to trigger mortality through the combination of hydraulic failure and carbon starvation (Adams et al. 2017; Anderegg et al. 2012; McDowell et al. 2011).

Another challenge when studying mortality is that the physiological processes governing tree vulnerability may vary in space and time. For instance, vulnerability may vary among individual trees within a population according to (i) the spatial heterogeneity in available resources, especially soil water (Nourtier et al. 2014); (ii) the heterogeneity in an individual tree’s life history, and in particular the effects of past stresses on tree morphology and anatomy (Vanoni et al. 2016); (iii) the inter-individual variation of physiological responses to stresses, which depends on ontogenic, plastic, and genetic effects controlling the expression of traits (Anderegg 2015a; Vitasse et al. 2009). Vulnerability may also vary through time for a given individual/population, not only because of temporal climatic variation but also through inter-individual variations in phenological processes. This is well illustrated by the risk of late frost damage, which is closely related to the coincidence between temporal patterns of budburst phenology, and the climatic sequence of low temperatures. Although relatively large safety margins were found regarding the risk of late frost damage during budburst across many European temperate tree (Bigler and Bugmann 2018), these safety margins may reduce with climate change, due to earlier budburst (Augspurger 2009). When young leaves have been damaged, some species can reflush, i.e. produce another cohort of leaves (Augspurger 2009; Menzel, Helm, and Zang 2015), but the time required to reflush may reduce the length of the growing season (Lenz et al. 2013), and may lead to mortality if trees do not have enough reserves to do this.

Available approaches to investigate the multiple drivers and processes underlying tree mortality can be classified into two broad categories: statistical, observational approaches versus process-based, mechanistic approaches. Statistical approaches use forest inventory data to test which tree characteristics (e.g. related to tree size and growth rate, biotic and abiotic environment, including management) affect mortality. By comparing species or populations over areas with large climatic variations, such studies have demonstrated the overall effect of drought severity on mortality, although usually explaining only a limited proportion of the inter-annual variance observed in mortality rate (Allen et al. 2010; Greenwood et al. 2017). Moreover, probabilities of mortality have been predicted with a higher accuracy when individual covariates for tree growth, size and/or competition were included in the statistical models, highlighting the importance of inter-individual variability in the threshold for mortality (Hülsmann, Bugmann, and Brang 2017; Monserud 1976). Recent statistical studies have attempted to include functional traits involved in the response to stress as additional covariates to improve the accuracy of mortality prediction, such as defoliation (Carnicer et al. 2011) or hydraulic safety margins (Benito-Garzón et al. 2018). Overall, the main advantage of statistical approaches is their ability to account for a potentially high number of factors and processes triggering mortality and for inter-individual variability in the threshold for mortality. However, these statistical models barely deal with the usually low temporal resolution of mortality data, missing information on the cause of tree death, and non-randomization inherent to natural population designs. In addition, the accuracy of statistical predictions can decrease outside the studied area (Hülsmann, Bugmann, and Brang 2017).

On the other hand, biophysical and ecophysiological process-based models (PBMs), initially developed to simulate carbon and water fluxes in forest ecosystems, are also useful to investigate the environmental drivers and physiological processes triggering tree mortality. For example, using the PBM CASTANEA, Davi & Cailleret (2017) showed that mortality of silver fir in southern France resulted from the combination of drought-related carbon depletion and pest attacks. Using six different PBMs, Mc Dowell et al. (2013) found that mortality depended more on the duration of hydraulic stress than on a specific physiological threshold. A main advantage of PBMs is their ability to understand how physiological processes drive mortality and to predict mortality under new combinations of forcing variables in a changing environment. However, they need a large number of parameters to be calibrated. Most often, calibration is made using the average parameters’ values known at species level, and therefore does not account for possible inter-individual variability of ecophysiological processes (Berzaghi et al. 2019). Moreover, biophysical and ecophysiological PBMs generally do not take into account individual tree characteristics (i.e. related to ontogenic, plastic and/or genetic variation). Hence, statistical and process-based approaches appear as complementary, and many authors have called for studies comparing or combining them (Hawkes 2000; O’Brien et al. 2017; Seidl et al. 2011).

In this study, we used both a statistical regression model and the PBM CASTANEA (Figure 1), to investigate the drivers of mortality within a population located at the warm and dry ecological margin for European beech (42° 28’ 41” N, 3° 1’ 26” E; Supplementary Figure S1). Mortality, decline (crown defoliation and fungi presence), size, growth, competition and budburst were characterised in a set of 4323 adult trees over a 15 year-period from 2002 to 2016. CASTANEA was used to simulate the number of late frost days, the percent loss of conductance (PLC) and the biomass of carbon reserves in response to stress. Specifically, we addressed the following questions: (1) How do climatic factors and physiological processes drive temporal variation in the mortality rate? (2) How do factors varying at tree-level modulate the individual tree’s probability of mortality? (3) How do physiological mechanisms modulate the vulnerability of individuals?

**Figure 1:**
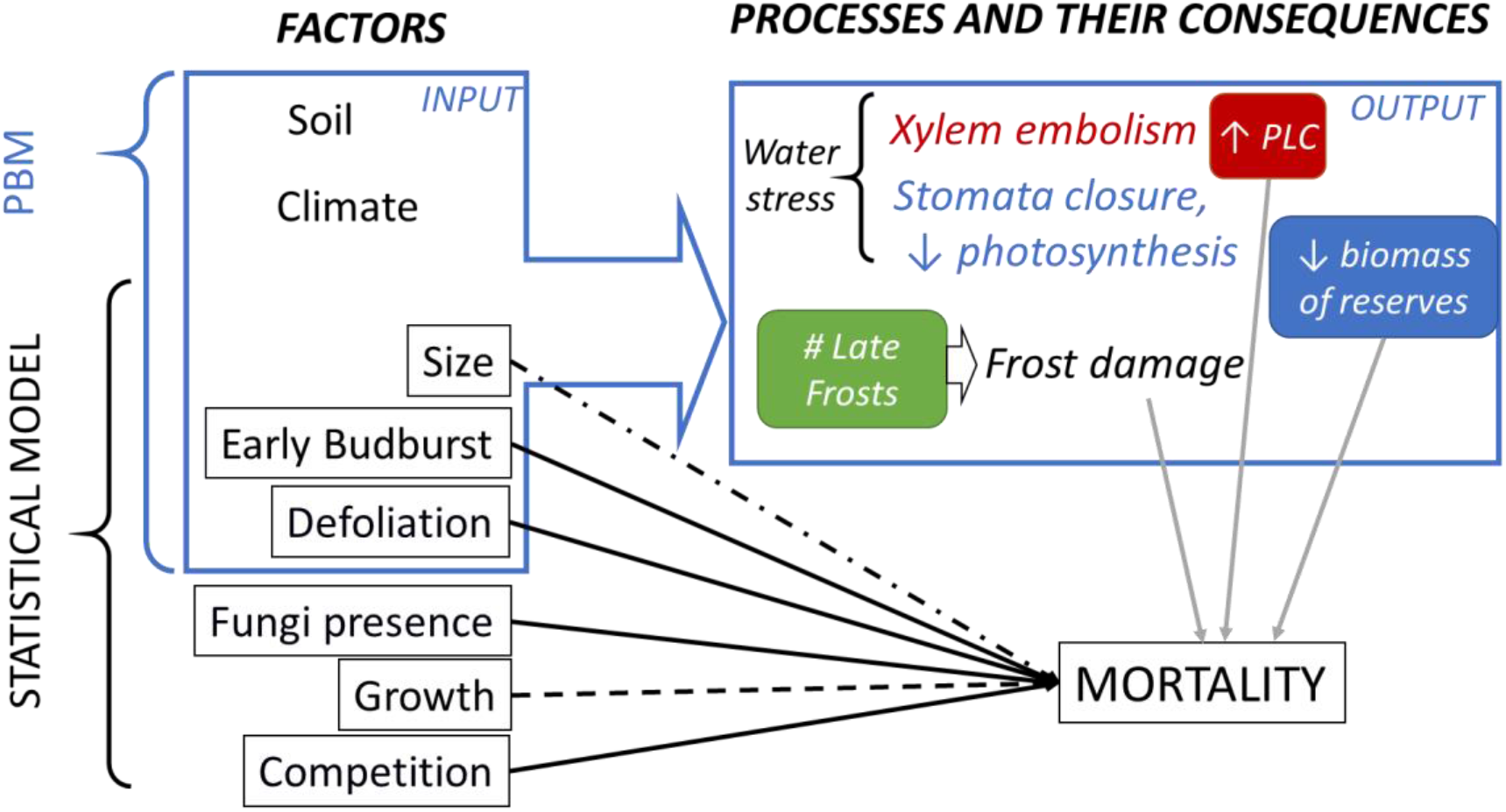
Combining process-based and statistical models to study variables and processes involved in tree mortality. The square boxes indicate the measured factors and response variables considered in statistical models. Boxes with rounded corners indicate stress-related output variables simulated with the PBM CASTANEA. The blue box on the left delineates the input variables of CASTANEA. At the top, grey arrows indicate the relationships considered to link stress-related output variables simulated by CASTANEA with observed mortality rate in the studied population. At the bottom, the black arrows indicate the relationships considered in the statistical model for the probability of mortality at individual level (solid lines: expected positive effect; dashed lines: expected negative effect; non-linear effects were expected for size). Moreover, the effects of size, early budburst and defoliation on the individual probability of mortality were also investigated using the PBM. PLC – percent loss of conductance.

## Materials and Methods

### Study species

The European beech (*Fagus sylvatica L.*) combines a widespread distribution (from northern Spain to southern Sweden and from England to Greece) and an expected high sensitivity to climate change (Cheaib et al. 2012; Kramer et al. 2010). Bioclimatic niche models predict a future reduction of this species at the rear edge of its range over the next few decades (Cheaib et al. 2012; Kramer et al. 2010). Its growth is highly sensitive to droughts (Dittmar, Zech, and Elling 2003; Jump, Hunt, and Penuelas 2006; Knutzen et al. 2017), which increase defoliation (Penuelas and Boada 2003). However, the low mortality rate observed so far in beech has led some authors to propose that this species presents a higher heat stress tolerance and metabolic plasticity when compared to other tree species (García-Plazaola et al. 2008). This apparent paradox between a low mortality and a high sensitivity to climate makes beech an interesting model species to study.

### Study site

La Massane is a forest of 336 ha located in the French eastern Pyrenees ranging from 600 to 1127 m.a.s.l. Located in the south of the beech range, the forest is at the junction of Mediterranean and mountainous climates with a mean annual rainfall of 1260 mm (ranging from 440 to 2000 mm) and mean annual temperature of 11°C (with daily temperature ranging from −10°C to 35°C) (Supplementary Figure S2). No logging operations have been allowed since 1886 and the forest was classified as a reserve in 1974. European beech is the dominant tree in the canopy representing about 68% of basal area of the forest. Beech is in mixture with downy oak (*Quercus pubescens* Willd), maple (*Acer opalus* Mill., *Acer campestris* L., *Acer monspessulanum* L.), and holly (*Ilex aquifolium* L.). A 10ha fenced plot has excluded grazing from livestock and large herbivores since 1956. All trees from this protected plot have been geo-referenced and individually monitored since 2002 (Supplementary Figure S3).

We estimated the soil water capacity (SWCa) through soil texture, soil depth and percentage of coarse elements measured in two soil pits in the protected plot. Secondly, we estimated the mean Leaf Area Index (LAI) by using hemispherical photographs (Canon 5D with Sigma 8mm EXDG fisheye). We computed the LAI and clumping index following the methodology described by Davi et al. (2009). SWCa and LAI were measured at population level.

### Individual tree measurements

This study is based on the characterisation of twelve variables in 4323 beech trees in the protected plot over the period from 2002 to 2016 (Table 1). Note that beech sometimes produces stump shoots resulting in multiple stems from a single position (coppice). Here about 10% of beeches occur in coppice, and each stem of all the coppices was individually monitored and subsequently referred as a “tree”.

**Table 1:** Quantitative (a) and categorical variables (b) measured at individual level. All the variables were measured in 4323 trees, except for H_2002_ (1199 trees). The “Cat” column indicates the category (i.e. size, growth, competition, decline, phenology). The “YMeas.” column indicates the year of measurement; note that all the variables were measured only once, so when two dates are given they indicate the period over which the variable is computed.

| Code | Variable | Cat | YMeas. | mean | min | max | unit |
| --- | --- | --- | --- | --- | --- | --- | --- |
| DBH<sub>2002</sub> | Diameter at breast height measured in 2002 | Size | 2002 | 21.7 | 10.0 | 116.0 | cm |
| DBH<sub>2012</sub> | Diameter at breast height measured in 2012 | Size | 2012 | 22.8 | 10.0 | 116.0 | cm |
| MBAI | Mean basal area increment between 2002 and 2012. | Growth | 2002-2012 | 4.7 | 0.0 | 95.0 | cm<sup>2</sup>. year<sup>-1</sup> |
| H<sub>2002</sub> | Height measured in 2002 | Size | 2002 | 8.8 | 2.0 | 26.0 | m |
| DEFw | Cumulated and weighted defoliation score | Decline | 2003-2016 | 0.1 | 0.0 | 1.0 | - |
| Nstem | Number of stems observed in the coppice | Compet | 2002 | 1.5 | 1.0 | 11.0 | - |
| Compet<sub>intra</sub> | Intra-specific competition index | Compet | 2002 | 2.7 | 0.0 | 11.4 | - |
| Compet<sub>intra+</sub> | Intra-specific competition index accounting for within-coppice competition | Compet | 2002 | 1.0 | 0.0 | 12.7 | - |
| Compet<sub>tot</sub> | Total competition index, accounting for within-coppice competition | Compet | 2002 | 4.6 | 0.1 | 20.0 | - |

(b)
| Code | Variable | Cat | YMeas. | Level | Number of trees |
| --- | --- | --- | --- | --- | --- |
| Fungi | Presence (1) or absence (0) of the saproxylic fungus | Decline | 2003-2016 | 1:<br>0: | 414<br>3913 |
| Budburst | Early (1) or late (0) budburst | Phenology | 2002 | 1:<br>0: | 237<br>4090 |

Tree mortality was recorded every year from 2003 to 2016, based on two observations (in autumn, based on defoliation and in spring, based on budburst). A tree was considered to have died at year *n* when (1) budburst occurred in the spring of year *n* but (2) no leaves remained in the autumn of year *n*, and (3) no budburst occurred in year *n*+1. All the 4323 trees were alive in year 2003 (Supplementary Figure S4). We computed the annual mortality rate (τ_n_) for each year *n* as:

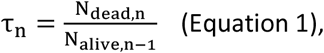

where *N_dead,n_* (respectively *N_alive,n_*) is the number of dead (respectively alive) trees in year *n*.

Diameter at breast height (DBH) was measured 1.30 m above ground level in 2002 and 2012. As we focused on the drivers of mature tree mortality, only trees with DBH_2002_ greater than 10 cm were retained for analysis. Individual growth was measured by the mean increment in basal area (MBAI) between 2002 and 2012, estimated as:

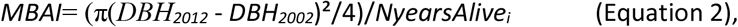

where *NyearsAlive_i_* is the number of years where individual *i* was observed being alive. Height in 2002 was estimated for a subset of 1199 trees.

A bimodal pattern in budburst phenology had been previously reported in La Massane (Gaussen 1958; Perci du Sert 1982). Some trees were observed to systematically initiate budburst about two weeks before all the others. Here, the monitoring allowed budburst phenology to be surveyed as a binary categorical variable, distinguishing trees with early budburst from the others.

The presence of defoliated major branches was recorded each year between 2003 and 2016 (except 2010) as a categorical measure (DEF = 1 for presence; DEF = 0 for absence). These annual measures were cumulated and weighted over the observation period for each individual in the following quantitative variable:

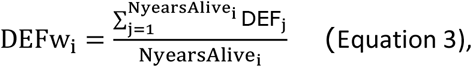

Year 2010 was not included in NyearsAlive_i_. DEFw integrates (without disentangling) the recurrence of defoliation and the ability to recover from defoliation. The presence of fructification of the saproxylic fungus *Oudemansiella mucida* (Schrad.) was recorded as a categorical measure (Fungi = 1 for presence; Fungi = 0 for absence). Given that once observed, the fructification persists throughout the subsequent years, we analysed it as a binary variable.

Competition around each focal beech stem was estimated by the number of stems in the coppice (*Nstem*) as an indicator of within-coppice competition. We also computed competition indices accounting simultaneously for the diameter (DBH_2002_) and the distance (d_ij_) of each competitor *j* to the competed individual *i*, following Martin and Ek (1984):

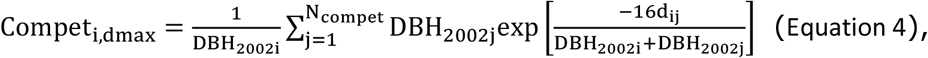

where *N_compet_* is the total number of competitors in a given radius dmax (in m) around each focal individual *i*. Only trees with DBH_2002_> DBH_2002i_ are considered as competitors. Such indices were shown to describe more accurately the competition than indices relying on diameter only (Stadt et al. 2007). We computed this competition index in three ways. The intra-specific competition index Compet_intra_ only accounts for the competition of beech stems not belonging to the coppice of the focal tree. The intra-specific competition index Compet_intra+_ accounts for all beech stems belonging, or not, to the coppice of the focal tree. The total competition index Compet_tot_ accounts for all stems and species. We considered that stems located less than 3 m away from the focal stem belonged to the same coppice. The three indices were first computed at all distances from 1 m (or 3 m for Compet_intra_) to 50 m from the target tree, with 1 m steps. We retained d_max_ = 15 m in subsequent analyses, because all indices plateaued beyond this threshold value, suggesting that in a radius greater than 15 m, the increasing number of competitors is compensated for by distance.

### Climate data

Local climate has been daily monitored *in situ* since 1976 and 1960 for temperature and precipitation/mean relative humidity, respectively. In order to obtain a complete climatic series (from 1959 to 2016), we used the quantile mapping and anomaly method in the R package “meteoland” (De Caceres et al. 2018), considering the 8-km-resolution-SAFRAN reanalysis (Vidal et al. 2010) as reference.

From the corrected climate series, we derived the daily climatic input variables for CASTANEA, which are the minimum, mean and maximum temperatures (in °C), the precipitation (mm), the wind speed (m.s^-1^), the mean relative humidity (%) and the global radiation (MJ.m^-2^).

### Simulations with CASTANEA

#### Model overview

CASTANEA is a PBM initially developed to simulate carbon and water fluxes in forest ecosystems with no spatial-explicit representation of trees (Dufrêne et al. 2005). A tree is abstracted as six functional elements: leaves, branches, stem, coarse roots, fine roots and reserves (corresponding to non-structural carbohydrates). The canopy is divided into five layers of leaves. Photosynthesis is half-hourly calculated for each canopy layer using the model of Farquhar et al. (1980), analytically coupled to the stomatal conductance model proposed by Ball et al. (1987). Maintenance respiration is calculated as proportional to the nitrogen content of the considered organs (Ryan 1991). Growth respiration is calculated from growth increment combined with a construction cost specific to the type of tissue (De Vries, Brunsting, and Van Laar 1974). Transpiration is hourly calculated using the Monteith (1965) equations. The dynamics of soil water content (SWCo; in mm) is calculated daily using a three-layer bucket model. Soil drought drives stomata closure via a linear decrease in the slope of the Ball et al. (1987) relationship, when relative SWCo is under 40% of field capacity (Granier, Biron, and Lemoine 2000; Sala and Tenhunen 1996). In the carbon allocation sub-model (Davi et al., 2009; Davi & Cailleret 2017), the allocation coefficients between compartments (fine roots, coarse roots, wood, leaf and reserves) are calculated daily depending on the sink force and the phenological constraints. CASTANEA model was originally developed and validated at stand-scale for beech (Davi et al. 2005).

#### Focal processes and output variables

In this study, we focussed on three response variables simulated by CASTANEA: (1) the percent loss of conductance (PLC) as an indicator of vulnerability to hydraulic failure; (2) the number of late frost days (NLF) as an indicator of vulnerability to frost damage; and (3) the biomass of reserves (BoR) as an indicator of vulnerability to carbon starvation. Note that we did not simulate mortality with CASTANEA because the thresholds in PLC, NLF and BoR triggering mortality are unknown. These variables were simulated using the CASTANEA version described in Davi and Cailleret (2017) with two major modifications. First, for budburst, we used the one-phase UniForc model, which describes the cumulative effect of forcing temperatures on bud development during the ecodormancy phase (Chuine, Cour, and Rousseau 1999; Gauzere et al. 2017). We simulated damage due to late frosts (see details in Appendix 1) and considered that trees were able to reflush after late frosts. We calculated NLF as the sum of late frost days experienced after budburst initiation.

Second, we implemented a new option in CASTANEA to compute PLC following the formula of Pammenter and Willigen (1998):

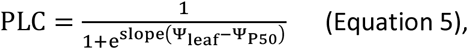

with Ψ_leaf_ (MPa) the simulated midday leaf water potential, Ψ_50_ (MPa) the species-specific potential below which 50% of the vessels are embolized, and *slope* a constant fixed to 50.

The leaf water potential Ψ_leaf_ was calculated as:

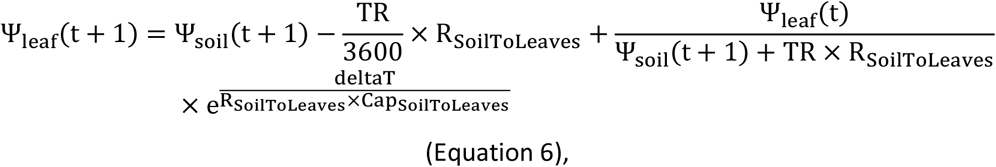

where the soil water potential (Ψ_soil_ MPa) was calculated from daily SWCo (Campbell 1974). Ψ_leaf_ was calculated hourly (deltaT = 3600s) based on the sap flow (TR in mmol.m^-2^.leaf^-1^) simulated following the soil-to-leaves hydraulic pathway model of Loustau et al. (1990). We used a single resistance (R_SoilToleaves_ in MPa.m^2^.s^1^.kg^-1^, following Campbell 1974) and a single capacitance (Cap_SoilToleaves_ in kg.m^-2^.MPa^-1^) along the pathway. R_SoilToleaves_ was assessed using midday and predawn water potentials found in the literature.

We added a binary option in CASTANEA to simulate branch mortality and defoliation as a function of PLC. In our case, when defoliation capability was added to the simulation with the option, we traduced the loss of leaves by reducing the LAI of the simulated tree. If the PLC at year *n* was >0, the LAI at year *n* was reduced by the PLC value for trees able to defoliate (option “Defoli-able”). Otherwise, PLC has no consequences on LAI (no defoliation possible).

#### Simulation design

The aim of the first simulations was to investigate whether response variables simulated by CASTANEA correlated with patterns of observed mortality in the studied population. We simulated a population of 100 trees representing the variability in individual characteristics observed in La Massane in terms of height-diameter allometry, DBH, leaf area index and budburst phenology (Appendix 1). We also simulated a range of environmental conditions representing the observed variability in SWCa and tree density. We also used this first simulation to validate CASTANEA based on the correlation between simulated and observed ring width (Appendix 1).

For this first simulation, the values of focal output variables (BoR, PLC and NLF) were averaged across the 100 trees. We also computed a cumulated vulnerability index (CVI) for each year *n* combining the simulated BoR, PLC and NLF as follows:

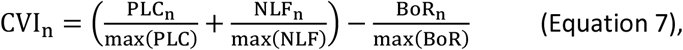

Note that each term is weighted by its maximal value across all years, so that the contribution of the three drivers to vulnerability is balanced. The possible range of CVI is [-1; 2].

The second set of simulations aimed at investigating the differences in physiological responses between individuals with different characteristics. We simulated eight individuals corresponding to a complete cross design with two size categories (5 and 40 cm in DBH), two budburst types (early and normal), and two defoliation levels (option “Defoli-able” activated or not).

### Statistical models of mortality to explore individual drivers of mortality

We used logistic regression models to investigate how tree characteristics affect the individual probability of mortality (P_mortality_). This approach is appropriate for a binary response variable and a mixture of categorical and quantitative explanatory variables, which are not necessarily normally distributed (Hosmer and Lemeshow 2000). We considered the following four full logistic regression models, which differ for the competition variable (separated below by “OR”):

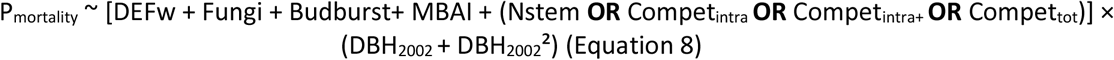

where the predictors defoliation (DEFw), growth (MBAI), size (DBH_2002_) and competition (Nstem or the Compet indices) were quantitative variables, and the presence of fungi (Fungi) and budburst phenology (Budburst) were categorical variables. We included both a linear and quadratic effect of DBH_2002_ by specifying this effect as a polynomial of second degree. Interaction effects of the previous predictors with this polynomial were included.

All variables were scaled before fitting the models. To select the best model depending on the choice of competition variable, we first fitted the full model described by equation 8 with each competition variable successively (Appendix 3). Then, we used the R package “MuMin” to compare and select the most parsimonious model among the four studied, based on AIC (Bartoń 2020).. Once de best competition variable chosen, our objective was to understand factors related to mortality rather than to achieve the best prediction, so we kept all the variables as recommended by Heinze et al. (2018) and Lederer et al. (2019). Model validity was checked based on the leverage points (i.e. points having a greater weight than expected by chance) with the Cook’s distance (Cook distance < 0.5 indicate no leverage). We evaluated the goodness-of-fit with the Brier test score (Brier 1950). We evaluated the sensitivity and specificity of the model using the receiver operating characteristic (ROC) curve.

Collinearity resulting from correlations between predictor variables is expected to affect the statistical significance of correlated variables by increasing type II errors (Schielzeth 2010). To evaluate this risk, we first checked for correlation among predictors included in equation 9 (Figure S5). We also computed the variation inflation factor (VIF) with the R package “car”. A threshold of the generalized VIF (GVIF) < 2 is commonly accepted to show that variables are not excessively correlated and do not make the model unstable.

We expressed the results in terms of odds ratios, indicating the degree of dependency between variables. For instance, the odds ratio for mortality as a function of budburst characteristics (early vs normal) is:

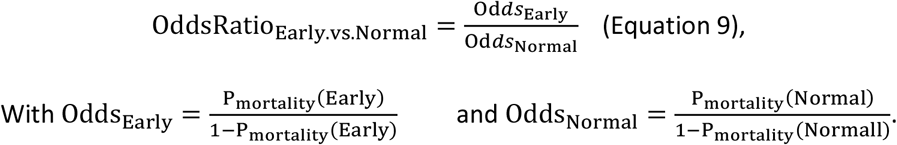

We computed odds ratios with “questionr” the R package (Barnier, Briatte, and Larmarange 2018). The interactions were visualized with the package “jtools” (Long 2018).

## Results

### Temporal variations in population mortality rate in relation to physiological vulnerability simulated with CASTANEA

We found a significant positive correlation between observed and simulated ring widths (p-value ≪ 0.01). Although CASTANEA tended to overestimate growth at the beginning of the simulated period, and simulated a decreasing trend in growth over time bigger than that in the observations. This is likely to be due to a bad estimation of population density prior to the monitoring period (see details in Appendix 1).

The cumulated mortality rate between 2004 and 2016 was 23% (Figure 2; Table S1). After 2004 (2.6%), two peaks of high annual mortality were observed, in 2006-2007 (3.3% in 2006) and in 2010 (2.9%). The lowest annual mortality rate was observed in 2008 (0.8%).

**Figure 2:**
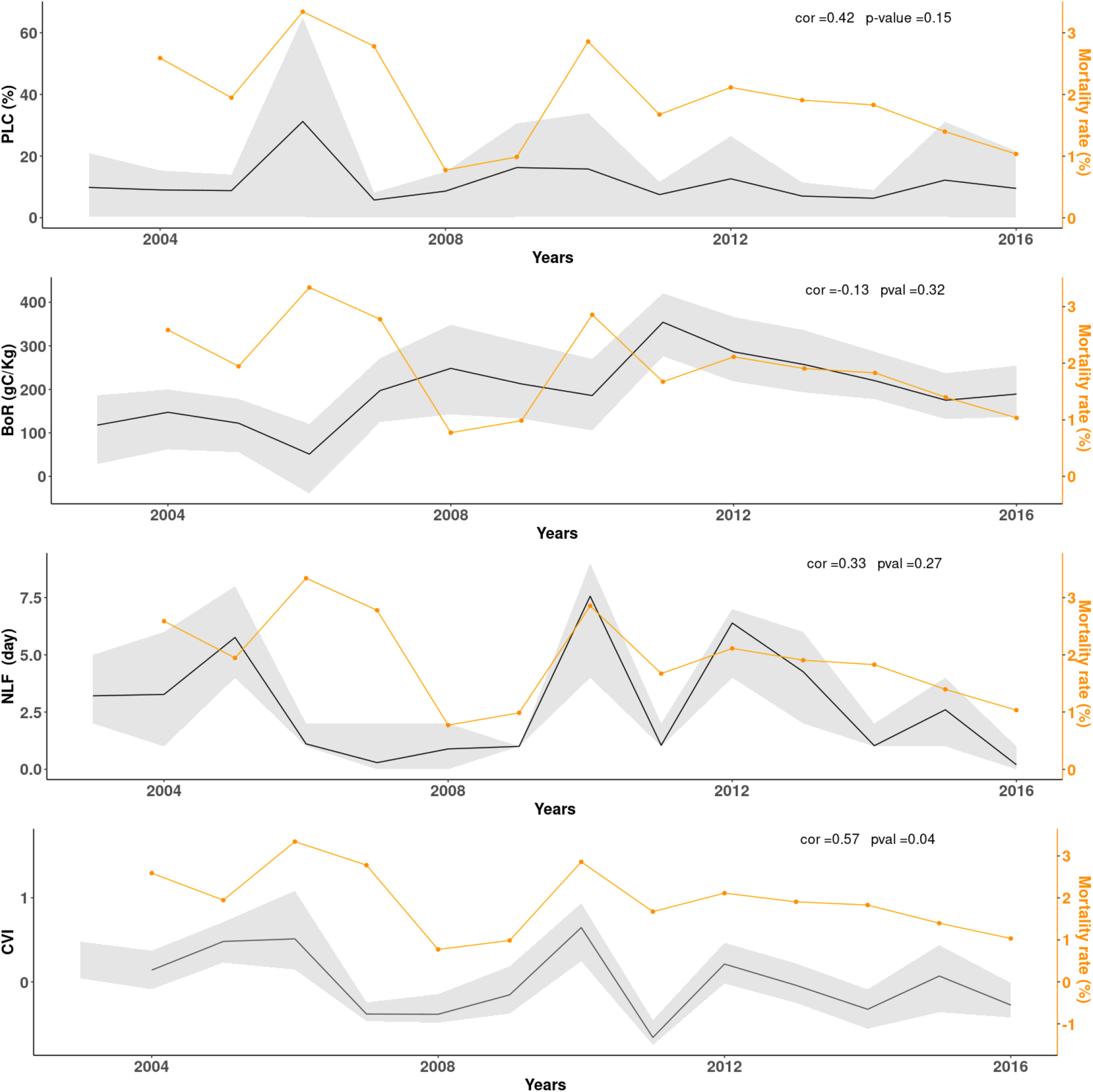
Stress-related output variable simulated with CASTANEA from 2004 to 2016: (a) percent loss of conductance (PLC); (b) biomass of reserves in gC.kg^-1^ (BoR); (c) number of late frost days (NLF); (d) Cumulated vulnerability index (CVI) integrating a, b and c. The black line is the mean of simulation, and the grey area represents the inter-individual variation from the 1^st^ to the 3^rd^ quartile. The yellow line is the mortality rate observed in La Massane.

CASTANEA simulated inter-annual variations in the percent loss of conductance (PLC): the mean PLC value varied among years, from 10% in 2004 and 2005 to 31% in 2006 (Figure 2a). The mean simulated biomass of carbon reserves (BoR) varied among years, from 51 gC.m^-2^ in 2006 to 354 gC.m^-2^ in 2011. Finally, the number of late frost days (NLF) varied among years, from 0.2 in 2016 to 7.56 days in 2010 (Figure 2c). The variation in the cumulative vulnerability index (CVI) integrated these different responses (Figure 2d), showed a peak in 2006 (drought), in 2010 (late frost) and in 2012 (combination of frost and drought).

None of the response variables simulated by CASTANEA (NLF, PLC, BoR) was alone significantly correlated to annual variation in mortality rate. However, a significant correlation was observed between CVI and the annual mortality rate (r = 0.58, p-value = 0.04). Hence, inter-annual variations in CVI were a good predictor of the mortality rate, except in year 2007. Besides the stress-related variables simulated with CASTANEA, we also investigated the effects of climatic variables on inter-annual variations in mortality rates using a beta-regression model (Appendix 2). We considered drought indices computed from climatic series, and this approach confirmed the effect of drought on mortality.

### Inter-individual variation in vulnerability simulated with CASTANEA

Simulations with CASTANEA showed that inter-individual differences in tree size, phenology, and defoliation, together with the intensity of climatic stress, affected the physiological responses to stress. The magnitude of the individual effects of each variable on tree vulnerability differed during a drought year (2006), a frost year (2010) and a good year (2008, 2014 or 2016; Figure 3). The loss of conductance was higher for trees with early budburst and for larger trees, but this effect was only evident in drought years (Figure 3a). Moreover, during drought, the ability to defoliate decreased the risk of cavitation (Figure 3a) but increased the risk of carbon starvation (Figure 3b). By contrast, phenology only poorly affected the biomass of reserve (BoR): even during a frost year, trees with earlier budburst did not reduce their BoR, due to their ability to reflush (Figure 3b). BoR was always lower for large tree, even without stress (Figure 3b). This was expected, because there is no explicit competition for light in CASTANEA. Hence large trees and small trees have a relatively similar photosynthesis when it is scaled by soil surface (large trees photosynthesise slightly more because they have a stronger LAI). Large trees, on the other hand, have a larger living biomass and thus a higher level of respiration, which leads to lower reserves (Table S2).

**Figure 3:**
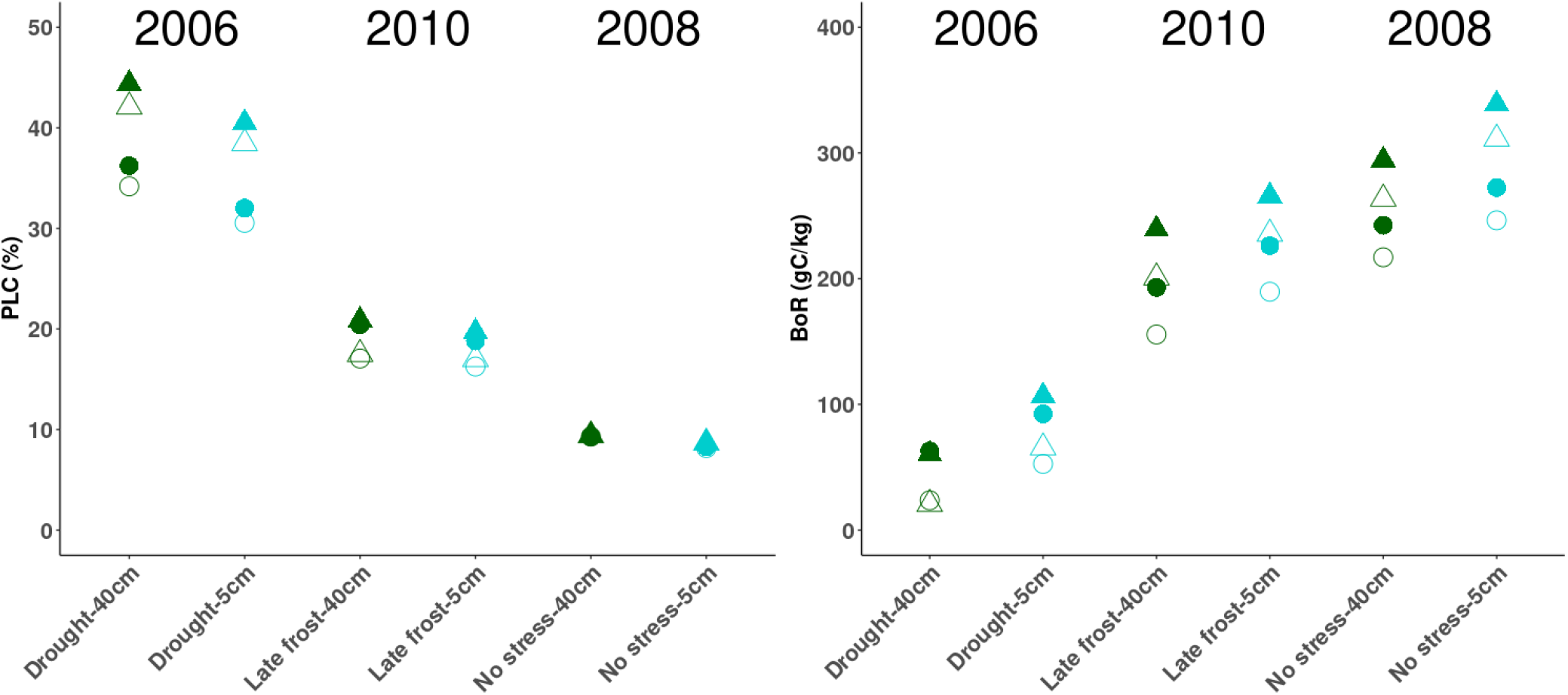
Physiological proxies of vulnerability simulated for eight trees differing in size, defoliation and budburst phenology. We focus on three key years: 2006 (drought); 2008 (no stress); 2010 (late frosts). Colours indicate the DBH at the beginning of the simulation: 5 cm (light blue) versus 40 cm (dark green). Triangles (respectively round) indicate individuals with early (respectively “normal”) budburst. Empty (respectively full) indicate individuals able (respectively not able) to defoliate. PLC: percentage of loss conductance; BoR: biomass of reserves.

### Inter-individual variation in the probability of mortality

Among the four models tested, the more parsimonious one was the one with the variable of competition N_stem_. All the variables listed in equation 8 had significant main effects on the probability of mortality, except DBH in its linear form (Table 2). This model explained 49% of the observed mortality and had both a good validity and goodness-of-fit (Appendix 3). However, the GVIF score for DBH was superior to 2 meaning that the collinearity with other variables was high, which we interpret as a consequence of the high number of interactions tested and not significant. Defoliation had the strongest linear effect on mortality: the relative probability of mortality increased by 1020 times for a one-unit increase in DEFw. Then, the relative probability of mortality was 2.26 higher for trees with earlier budburst as compared to others, and 1.88 higher for trees bearing fungi fructifications as compared to others. The relative probability of mortality increased with increasing N_stem_, and decreased with increasing MBAI. Regarding the effect of tree size, the polynomial of degree 2 corresponded to a U-shape and traduced a higher relative probability of mortality for both the smaller and the larger trees (In addition, this calibration is based on the mean of the individuals’ 2, Appendix 3).

**Table 2:** Effects of tree characteristics on the individual tree’s probability of mortality. Variables are defined in Table 1. Effects were estimated with a logistic regression model (equation 8). β is the maximum likelihood estimate, with its estimated error (SE), z-value, and associated p-value. OR is the odds ratio.

| Variables | $\beta$ | SE | z value | p-value | OR |
| --- | --- | --- | --- | --- | --- |
| DEFw | 6.93 | 0.26 | 26.30 | <0.0001 | $1.02 \cdot 10^3$ |
| Fungi | 0.63 | 0.16 | 3.96 | <0.0001 | 1.88 |
| Budburst | 0.82 | 0.17 | 4.71 | <0.0001 | 2.26 |
| MBAI | -0.45 | 0.08 | -5.58 | <0.0001 | 0.64 |
| Nstem | 0.13 | 0.04 | 3.44 | <0.0001 | 1.14 |
| DBH<sub>2002</sub> | -9.19 | 8.16 | -1.13 | 0.26 | $1.02 \cdot 10^{-4}$ |
| DBH<sub>2002</sub><sup>2</sup> | 21.32 | 9.32 | 2.29 | 0.02 | $1.81 \cdot 10^9$ |
| DEFw:DBH<sub>2002</sub> | -49.71 | 15.51 | -3.20 | $5.91 \cdot 10^{-4}$ | $2.57 \cdot 10^{-22}$ |
| DEFw:DBH<sub>2002</sub><sup>2</sup> | 32.97 | 17.08 | 1.93 | 0.05 | $2.08 \cdot 10^{14}$ |
| Fungi:DBH<sub>2002</sub> | -11.33 | 8.69 | -1.30 | 0.19 | $1.21 \cdot 10^{-5}$ |
| Fungi:DBH<sub>2002</sub><sup>2</sup> | -0.44 | 8.51 | -0.05 | 0.96 | 0.65 |
| Budburst:DBH<sub>2002</sub> | 0.27 | 10.41 | 0.03 | 0.98 | 1.31 |
| Budburst:DBH<sub>2002</sub><sup>2</sup> | -3.11 | 12.19 | -0.26 | 0.80 | $4.46 \cdot 10^{-2}$ |
| MBAI:DBH<sub>2002</sub> | 2.89 | 3.37 | 0.86 | 0.39 | 18.10 |
| MBAI:DBH<sub>2002</sub><sup>2</sup> | -6.18 | 3.34 | -1.85 | 0.06 | $2.07 \cdot 10^{-3}$ |
| Nstem:DBH<sub>2002</sub> | 1.32 | 4.12 | 0.32 | 0.75 | 3.76 |
| Nstem:DBH<sub>2002</sub><sup>2</sup> | 0.00 | 4.88 | 0.00 | 1.00 | 1.00 |

Interaction effects between diameter and defoliation on mortality were significant: the relative probability of mortality increased more rapidly with DEFw for small rather than larger trees, and at an equal level of defoliation, the probability of mortality was always higher for smaller trees (Figure 4a, Table 2). Interaction effects between DBH_2002_^2^ and growth on mortality were also significant: the decrease in the relative probability of mortality with increasing mean growth was evident mostly for small trees (Figure 4b). These results were robust whatever the choice of the competition variable (N_stem_ versus competition indices), or the choice of the size variable (height instead of diameter) and the choice of considering size (DBH_2002_) as a quantitative or a categorical variable (Appendix 3). Finally, we obtained similar results with an alternative approach (survival analysis) which account simultaneously for both levels of variability (individual and temporal) in our data set (Appendix 4).

**Figure 4:**
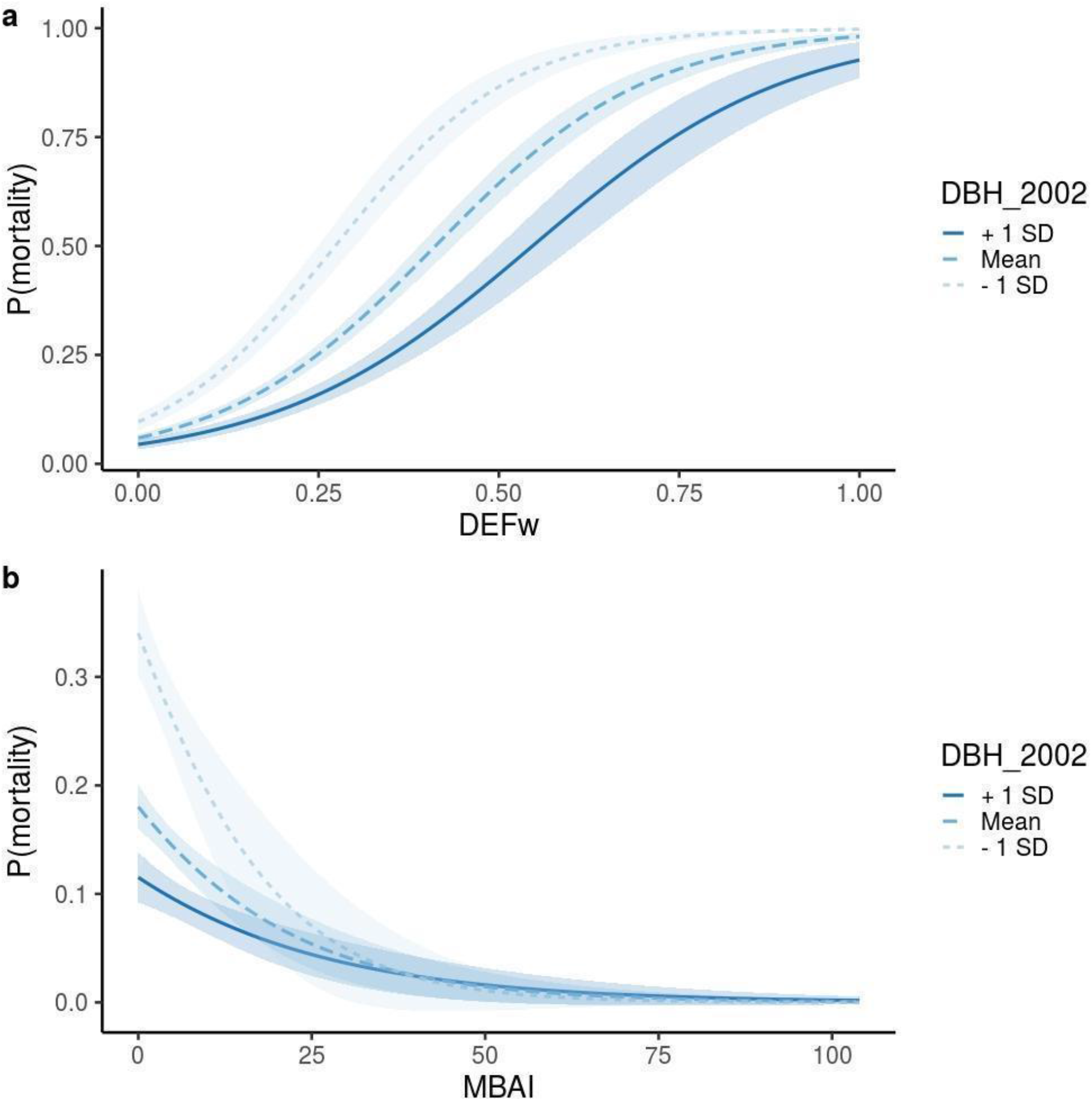
Interaction effects in the logistic regression model for individual mortality rates. (a) between diameter (DBH_2002_) and weighted defoliation (DEFw). (b) between DBH_2002_ and the mean growth in basal area (MBAI). Regression lines are plotted for three values of DBH_2002_, corresponding to ± 1 standard deviation (10.7 cm) from the mean (22 cm). Confidence intervals at 95% are shown around each regression line.

## Discussion

By comparing statistical and process-based models, this study shed new light on the inter-annual and inter-individual variability of mortality in a drought- and frost-prone beech population. We showed that mortality in this marginal population is triggered by a combination of climatic factors, and that the vulnerability to drought and frost is modulated by several individual characteristics (defoliation, vegetative phenology, growth, size, competitors), as summarized in Table 3.

**Table 3:**
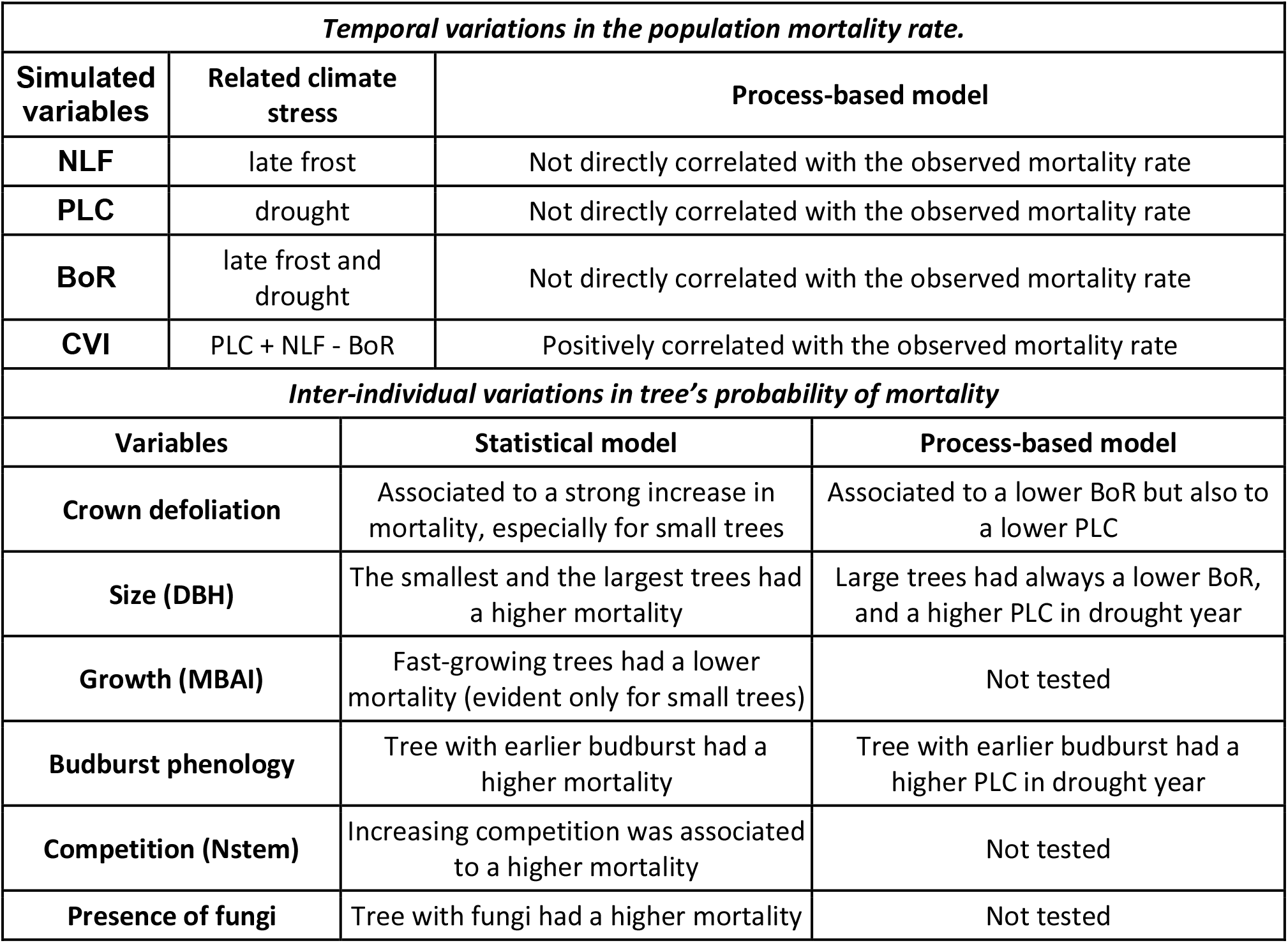
Summary of the main effects of the studied variables on mortality. NLF: number of late frost days; PLC= percentage of loss conductance; BoR: Biomass of reserve

### The rate of mortality increased in response to drought and late frosts

The annual mortality rates observed in this study ranged between 0.7 and 3.3% (mean value = 2%). This is at the upper range of the few mortality estimates available for beech. Hülsmann et al. (2016) reported annual mean rates of mortality of 1.4%, 0.7% and 1.5% in unmanaged forests of Switzerland, Germany and Ukraine, with a maximum mortality rate of 2.2%. Archambeau et al. (2019) estimated even lower mortality rates (mean annual value = 3.8 10^-3^%, range = 3.7 10^-3^% to 3.8 10^-3^%) from European forest inventory data (including managed and unmanaged forests). Overall, these mortality rates are low when compared to other tree species; for instance, according to the French national forest inventory, the average mortality is 0.1% for beech against 0.3% on average for other species and 0.4% for spruce or 0.2% for silver fir (IFN 2016). The relatively high value of natural mortality observed here may result from the absence of management (which resulted in high tree density), combined with the population location being at the dry, warm margin of species distribution (Figure S1), where most population extinctions are expected in Europe (Thuiller et al. 2005). However, we cannot rule out that the size threshold in inventories, which differ between studies, also affects these different mortality estimates (e.g., a higher frequency of smaller trees increases the mortality rate).

We showed that inter-annual variations in the observed mortality rate in our studied population were significantly associated with variations in the cumulative vulnerability index (CVI) integrating the number of late frost days (NLF), the percent loss of conductance (PLC) and the biomass of carbon reserves (BoR) simulated by CASTANEA. We found no correlation when the three response variables simulated by CASTANEA were considered separately, highlighting that patterns of mortality in beech are driven by a combination of drought and late-frost stresses. In particular, simulations showed that in 2010 (a year without drought), the high mortality rate coincided with an extreme late frost event. This is consistent with the study of Vanoni et al (2016), which showed that both drought and frost could contribute to beech mortality. Our results also support the emerging consensus that mortality at dry, warm margins is not due either to carbon starvation or hydraulic failure, but is rather the result of a balance of all these responses (e.g. McDowell et al. 2011; Sevanto et al. 2014).

In future developments, the CVI could be refined in several ways. Its different components could be weighted based on ecophysiological knowledge. The CVI could also benefit from taking into account the temporal dynamics of mortality, such as the existence of positive or negative post-effects across years. The number of years of observations in this study did not allow to account for these lagged effects, which probably explains why the CVI failed to predict the high mortality observed in 2007. Indeed, the high mortality in 2007 was probably due to the lagged effect of the 2006 drought. Such lags between the weakening of a tree and its final death were shown for beech in Vanoni (2016) and silver fir in Davi & Cailleret (2017).

### The vulnerability to drought and frost varied among individuals

The large number of trees individually monitored each year provided us with an exceptionally large sample size to test the inter-individual vulnerability to climatic hazards (drought and late frost) and to biotic pressures (competition and the presence of a fungus). Firstly, we found that a higher mean growth was associated with a lower probability of mortality, as previously demonstrated (Cailleret and Davi 2011; Gao et al. 2018). This decrease in mortality with increasing mean growth was evident mostly for small trees as already reported in beech seedlings (Collet and Le Moguedec 2007) and other species (Kneeshaw et al. 2006; Lines, Coomes, and Purves 2010), but not in adult beech trees to our knowledge.

Secondly, we found that increased defoliation was associated with increased mortality. This result was expected from previous studies (Dobbertin and Brang 2001, Carnicer et al. 2011), although the consequences of defoliation are still being debated for beech. Senf et al. (2018) showed that defoliation was associated with tree decline, while Bauch et al., (1996) and Pretzsch (1996) found that the growth of highly defoliated beech trees did not decrease and could even increase in some cases. Our simulations comparing trees able, or not, to defoliate, shed light on the multiple effects of defoliation on mortality. These simulations showed that defoliation decreased carbon reserves in good years but could also limit the loss of hydraulic conductance during dry years. Furthermore, we observed a significant interaction between defoliation and tree size on mortality, showing that small trees were more vulnerable to mortality in response to defoliation than large trees. However, we cannot rule out that this effect is due in part to the categorical method used to survey defoliation, which does not take into account the percentage of crown loss. Hence, defoliation may be biased with respect to size, such that small and defoliated trees will on average have a higher proportion of canopy loss, and therefore be more impacted than large and defoliated trees.

Thirdly, both statistical and process-based approaches found that trees with early budburst were more prone to die. By contrast, Robson et al. (2013) showed that trees with early budburst were not more vulnerable to mortality, but rather grew better, consistent with our simulations where trees with early budburst accumulate more reserves during good years. This discrepancy may be due to the location of our studied population at the rear-edge of beech distribution, where earlier budburst dates may expose trees to a higher risk of late frost. We can hypothesize that early budburst trees have been maintained in this population until now because they grow better in the “good” years, and therefore, are more likely to become dominant tree and have abundant reproduction. In CASTANEA simulations, the higher vulnerability of early trees resulted rather from a higher risk of hydraulic failure than from a higher impact of late frosts. This is because trees with early budburst have a longer vegetation season and they develop their canopies faster, which also increases their water needs due to the increase of transpiration. Altogether, the relationships between phenology and mortality deserve further investigation, especially since the spatio-temporal variation of budburst patterns under climate change may produce complex spatio-temporal patterns of stresses (Vanoni et al. 2016).

Regarding the effect of size, the results differed between the statistical approach, where large trees died less than small ones, and the simulations, which predicted a greater vulnerability to drought of large trees. There may be several explanations for this discrepancy. The first reason is that CASTANEA simulates an average tree without explicit competition for light and water; hence not accounting for the higher observed background mortality in small trees as compared to large ones. In addition, CASTANEA also does not account for individual dominance status, which can affect the current carbon balance of a tree and hence its capacity to mitigate stress. In the studied population, large trees are more likely to be dominant, with better access to light resources promoting carbon accumulation, as compared to small trees, which are more likely to be suppressed. Another reason is that tree size may vary with environmental factors in the studied population, such that large trees have a tendency to occur on better soils. Therefore, the size effect observed through the statistical approach may reveal the confounding effect of spatial soil heterogeneity, not taken into account in the PBM. A measurement of water availability at individual tree level would be necessary to address this issue but was out of the scope of this study.

### Comparing statistical and process-based approaches allow to identify the causes of tree vulnerability

These two approaches illustrate the classical compromise between a fine understanding of physiological mechanisms driving mortality, with complex and expensive PBMs, versus efficient precision in local mortality predictions, with statistical models requiring less data, but having a weaker ability to generalize proximal causes. Most often, studies adopt either of the two approaches, and generally statistical approaches prevail (Hülsmann et al. 2016; Seidl et al. 2011). However, the two approaches are highly complementary, and combining them allows to decipher the respective roles of the drivers and mechanisms underlying tree mortality and to understand their variability among individuals or years (Hawkes 2000; O’Brien et al. 2017; Seidl et al. 2011). The two approaches can be compared at the individual level, as this study does, or they can be combined, as when we analysed the correlation between the observed mortality rate and simulated stress response variables. An upper level of integration would be inverse modelling, where observed mortality rates could be used to infer the physiological thresholds (e.g. in BoR, PLC and NLF) likely to trigger mortality (Davi & Cailleret 2017; Cailleret et al., 2020).

This study illustrated a classical difficulty in combining statistical and process-based approaches, related to the difference between observed variables and PBM parameters. For instance, the comparison of defoliated and non-defoliated trees does not have exactly the same meaning when using CASTANEA and the statistical approach. In CASTANEA, we compared trees, able versus unable to defoliate, while these average trees shared on average the same edaphic conditions. In the statistical approach, we compared trees with different levels of defoliation, but which also probably did not share the same edaphic and biotic conditions. Defoliation was thus also likely to be an indicator of the fertility of the environment, such that on shallow soils, defoliation was stronger and the probability of mortality increased. Hence, the correlation does not necessarily involve a causal relationship between defoliation and mortality.

The major benefit of our approach combining different approaches (statistical, process-based) at different scales (forest stand, individual) is to allow to relate the ecological patterns observed at an upper scale (forest stand, multi-year period) with the patterns observed at a lower scale where processes operate (individual, year). This ability to aggregate/disaggregate patterns is acknowledged as a powerful approach to understand apparent contradictions between patterns observed at different scales (Clark et al. 2011). There are however some limitations to the approaches we used here. First, none of them could fully account for the non-independence of climatic effects on mortality between years. Indeed, the effect of climatic variables at a given year may depend on other variables expressed in previous years. This was observed in beech, where several drought years finally led to a growth decline (Jump et al. 2006; Knutzen et al. 2017; Vanoni et al. 2016) or a modification in sap flow (Hesse et al. 2019). Moreover, the processes driving mortality may change through time as the most sensitive individuals are progressively eliminated, and/or the surviving trees become less and less sensitive (i.e. acclimation Niinemets 2010). Finally, the statistical model at the individual level could not fully make use of the repeated measurements of mortality over the years, partly because other individual variables were measured only once over the study period (except defoliation). Survival analyses could unfortunately not fully address this limitation (Appendix 4), and the development of a finely tuned Bayesian approach was out of the scope of this study. Besides methodological improvements, another extension to the present study would be to combine statistical and process-based approaches at a larger spatial scale, among populations across climatic gradients. This would allow the investigation of whether the respective drought and late frost sensitivity differ between the rear, core and leading edge of species distribution, as suggested by Cavin and Jump (2017).

## Data accessibility

The data set analysed in this preprint is available online under the zenodo repository (https://doi.org/10.5281/zenodo.3519315). Raw data can be obtained from JG, JAM and CH.

## Supplementary material

The process-based model CASTANEA is an open-source software available on capsis website: http://capsis.cirad.fr/

Supplementary materials (Figures and Tables) for this preprint are available on bioRxiv (645747).

## Author Contributions

JAM, JG, CH and EM measured and mapped all the trees. CPC performed the wood core analyses. CPC, FL and SOM designed and ran the statistical models. CPC and HD ran the PBM. CPC drafted the manuscript, and all authors contributed to its improvement.

## Acknowledgments

We are grateful to M. Cailleret, B. Fady, and N. Martin Saint Paul for discussions and comments on a previous version of this manuscript. We thank E.Walker and F.Bonneu for statistical discussions and advices, N. Mariotte for wood core sampling, and F. Guibal for their analyses. SOM and HD were funded by the EU ERA-NET BiodivERsA projects TIPTREE (BiodivERsA2-2012-15) and the ANR project MeCC (ANR-13-ADAP-0006). CP received funding from the European Union’s Horizon 2020 research and innovation programme under grant agreement No. 676876 (GenTree). Version 7 of this preprint has been peer-reviewed and recommended by Peer Community In Ecology (https://doi.org/10.24072/pci.ecology.100070).

## Conflict of interest disclosure

The authors of this preprint declare that they have no financial conflict of interest with the content of this article. SOM is one of the PCIEcology recommenders.

## Appendices

Four supplementary appendices are available on bioRxiv (645747):

Appendix 1: CASTANEA model, calibration and simulation design

Appendix 2: Beta-regression model for the temporal variations in the rate of mortality in the studied population

Appendix 3: Logistic regression models for the probability of mortality at tree-level

Appendix 4: Survival analysis the probability of mortality at tree- and year-levels

## Notes

https://doi.org/10.5281/zenodo.3519315

### Competing Interest Statement

The authors have declared no competing interest.

### Summary of Updates

This is version 7 of the manuscript, peer-reviewed and recommended by PCI Ecology (2020).

