## Supplementary material for "Comparing statistical and mechanistic models to identify the drivers of mortality within a rear-edge beech population": Suplementary figures and tables

### Supplementary Figures and Tables for the manuscript "Comparing statistical and mechanistic models to identify the drivers of mortality within a rear-edge beech population"

by Cathleen Petit-Cailleux, Hendrik Davi, François Lefevre, Christophe Hurson, Joseph Garrigue, Jean-André Magdalou, Elodie Magnanou and Sylvie Oddou-Muratorio

**Figure S1: European climatic niche of European beech as depicted by the sum of precipitation and the mean annual temperature.**

For the 1358 locations (black empty dots) where beech is documented as present by von Wühlisch (2008), we used climate series available from Weedon et al. (2014) to compute average values of each climatic variable from 1976 to 2008**.** The red full dot is the studied site of La Massane.

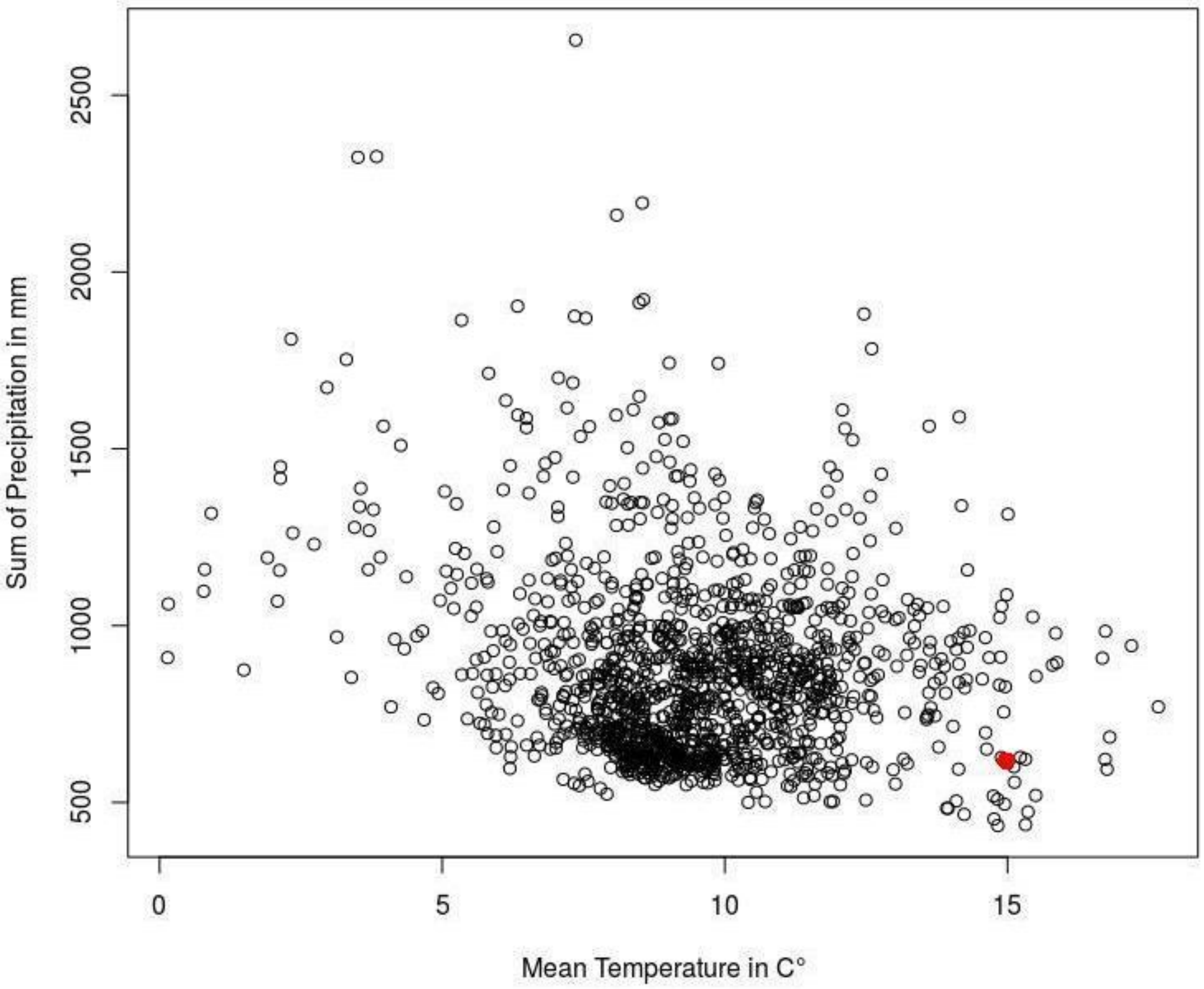

**Georg von Wühlisch**. **2008**. *EUFORGEN Technical Guidelines for genetic conservation and use for European beech ( Fagus sylvatica ).* Rome, Italy.

**Weedon GP, Balsamo G, Bellouin N, Gomes S, Best MJ, Viterbo P**. **2014**. The WFDEI meteorological forcing data set: WATCH Forcing Data methodology applied to ERA-Interim reanalysis data. *Water Resources Research* **50**: 7505–7514.

#### Figure S2: Climograph of the studied site (La Massane) between 1976 and 2016.

One graduation of the precipitation scale (in mm, on the rigth) corresponds to two graduations of the temperature scale (in °C, on the left) (P = 2T). Blue vertical lines indicate wet period. Dotted red vertical lines indicate drought period (P < 2T). Above the black horizontal lines indicate, monthly precipitation is greater than 100 mm, and the precipitation the scale is increased from 2mm/ºC to 20mm/ºC (P = 20T). Daily maximum average temperature of the hottest month and daily minimum average temperature of the coldest month are labelled in black at the left margin of the diagram. Light square blue indicates the probable frost months.

**Jose A. Guijarro (2019)**, Climate Tools (Series Homogenization and Derived Products), “climatol” package.

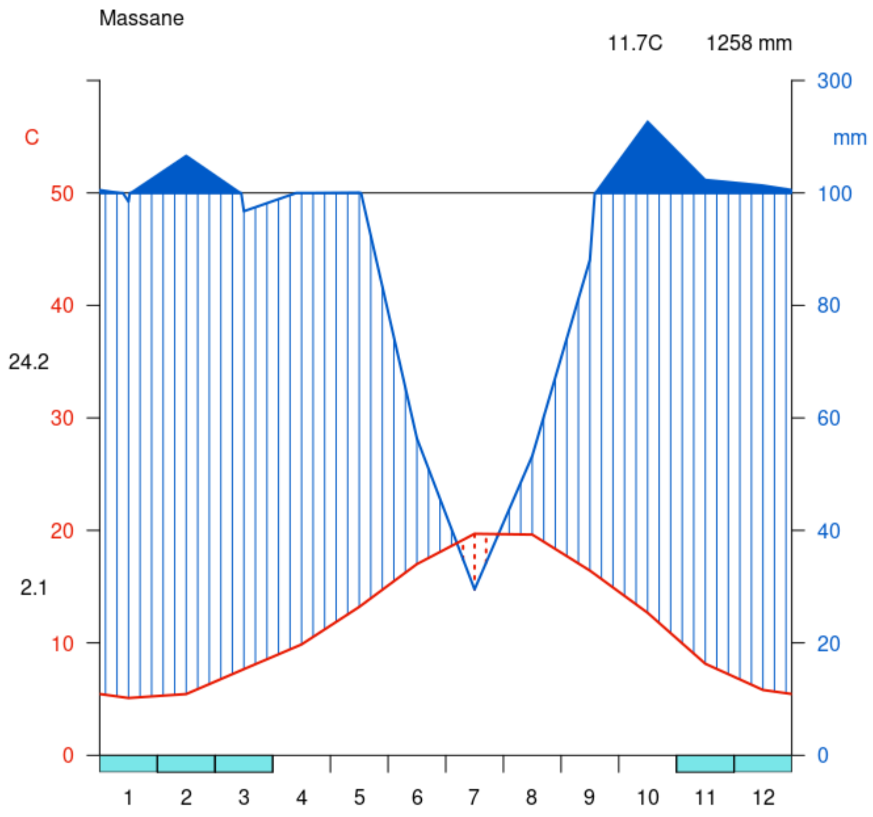

**Figure S3: Spatial distribution of trees in the studied site La Massane.**

Dot's size is proportional to tree diameter at breast height ($DBH_{2002}$) and only tree with DBH2002> superior to 10 cm are shown (and studied).

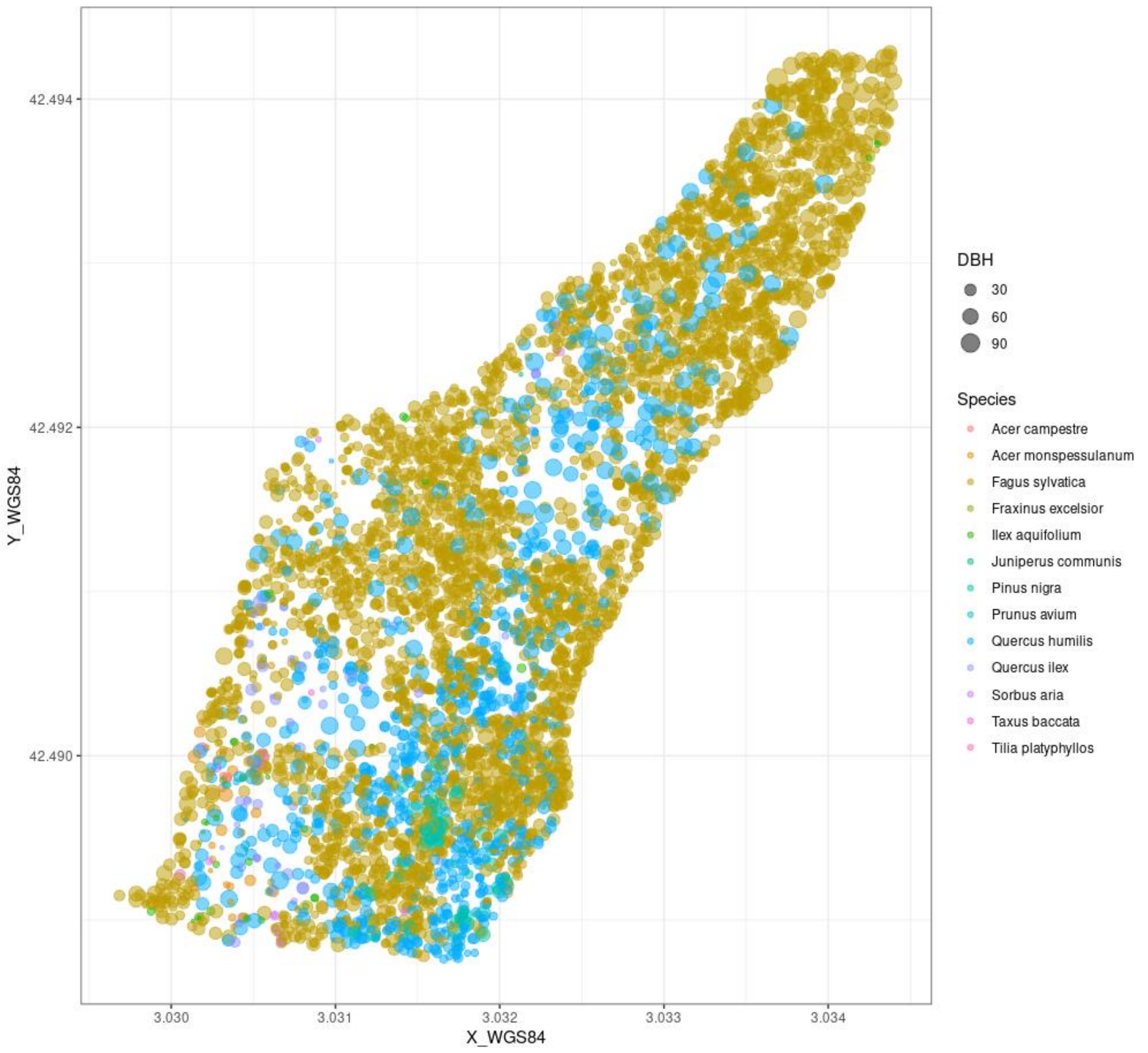

**Figure S4: Spatial distribution of alive (blue dots) and dead (pink dots) beech trees at the end of the studied period (2004-2016).**

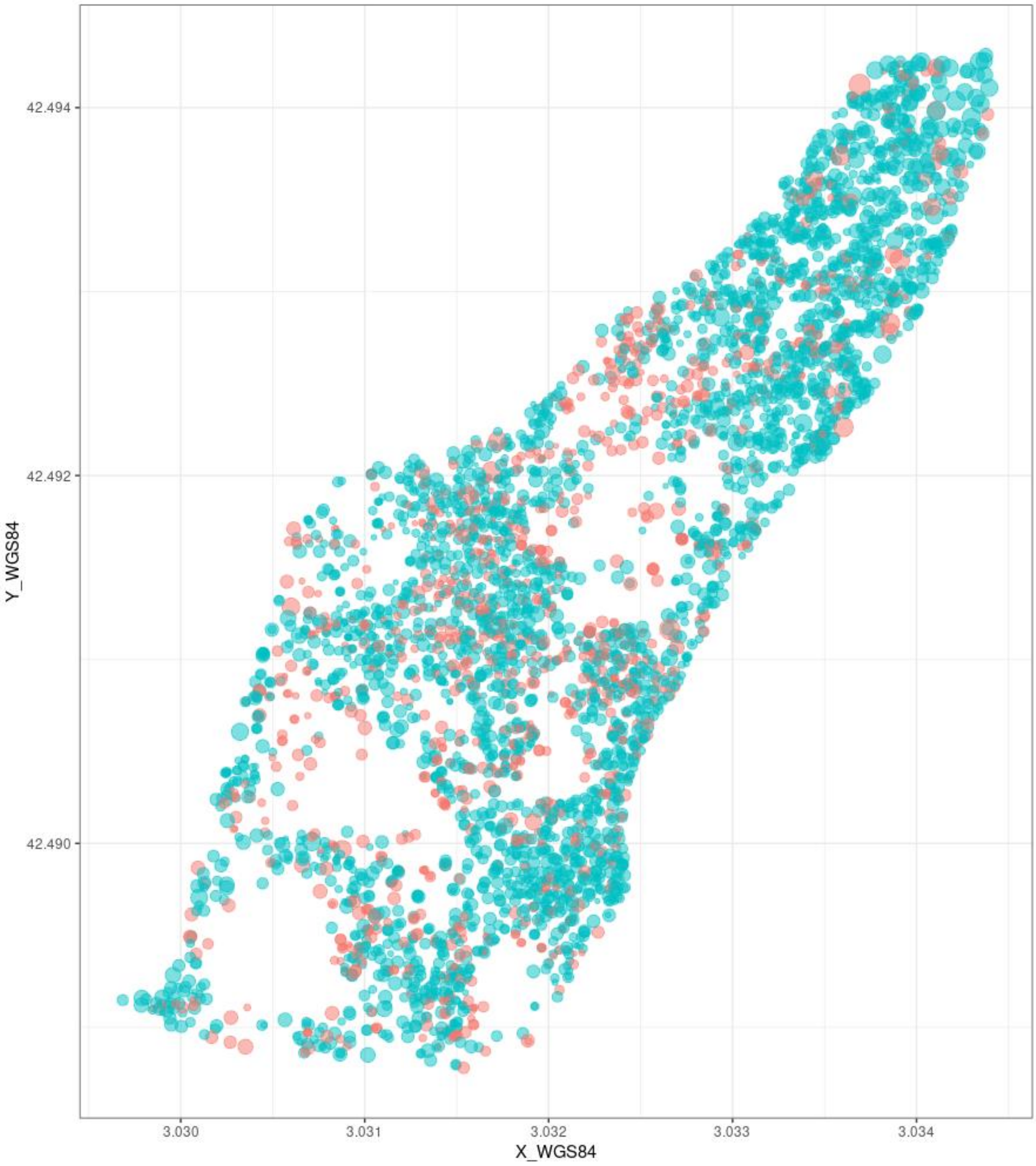

#### Figure S5: Pairwise distribution and correlations among quantitative variables

The names and meaning of variables are given in table 1.

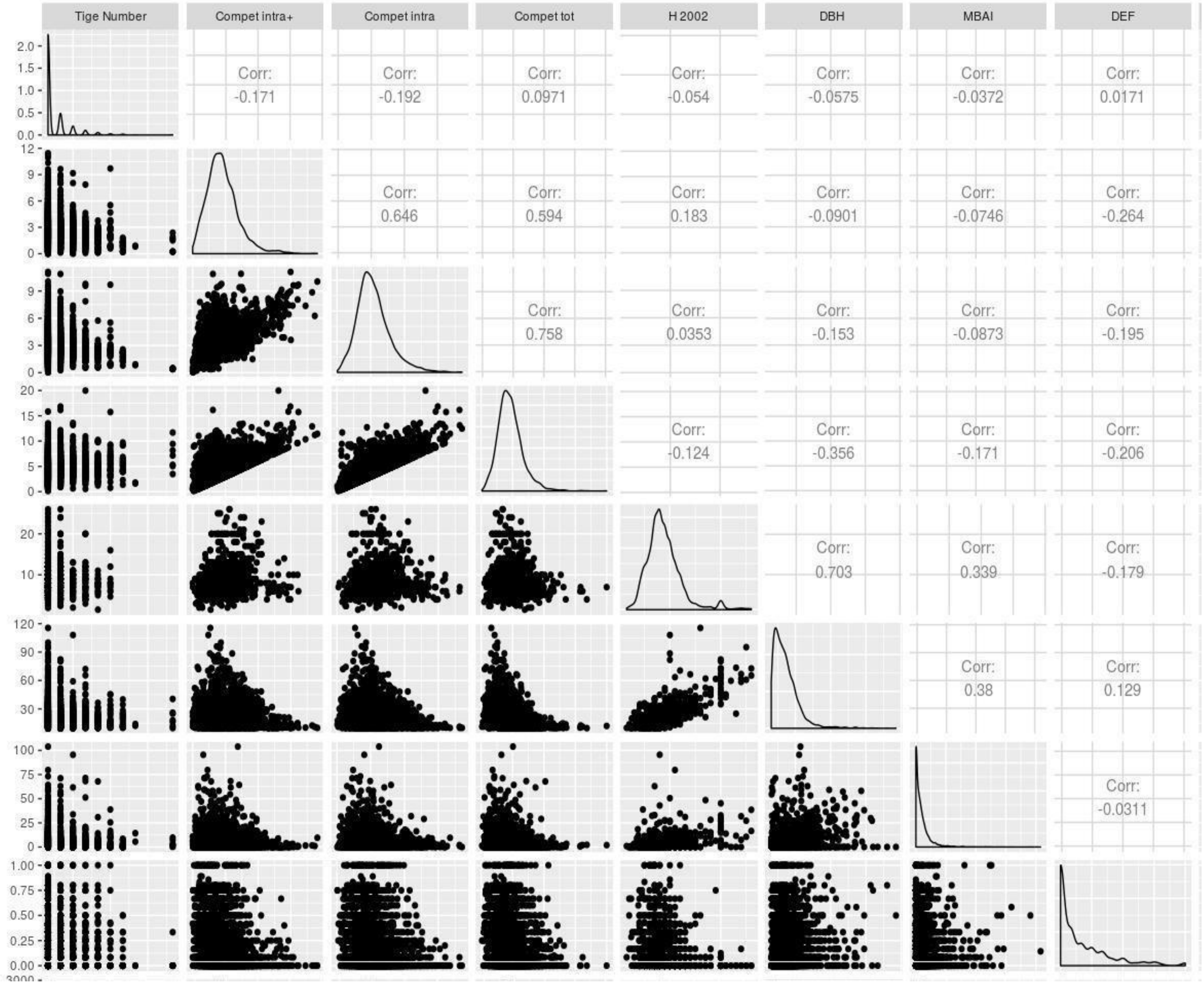

**Table S1 - Observed mortality rates and simulated stress-related output variables of CASTANEA from year 2004 to year 2016.**

| Year | 2004 | 2005 | 2006 | 2007 | 2008 | 2009 | 2010 | 2011 | 2012 | 2013 | 2014 | 2015 | 2016 |
|---|---|---|---|---|---|---|---|---|---|---|---|---|---|
| Mortality rate in % | 2.6 | 1.9 | 3.3 | 2.8 | 0.8 | 1.0 | 2.9 | 1.7 | 2.1 | 1.9 | 1.8 | 1.4 | 1.0 |
| PLC | 0.09 | 0.09 | 0.31 | 0.06 | 0.09 | 0.16 | 0.16 | 0.08 | 0.13 | 0.07 | 0.06 | 0.12 | 0.1 |
| BoR | 147.32 | 122.12 | 51.1 | 197.17 | 248.22 | 212.43 | 185.71 | 354.24 | 286.18 | 256.9 | 219.7 | 175 | 18912 |
| NLF | 3.27 | 5.76 | 1.11 | 0.29 | 0.89 | 1.00 | 7.56 | 1.05 | 6.39 | 4.26 | 1.03 | 2.6 | 0.2 |

**Table S2 –Mean and standard deviation values of output variables simulated by CASTANEA.**

The output variables of interest are: the Net Primary Production (NPP in gC.cm$^{-2}$. Year$^{-1}$), the Gross Primary Production (GPP in gC.cm$^{-2}$. Year$^{-1}$), Leaf Area Index (LAI) and autotrophic respiration (Rauto in gC.cm$^{-2}$. Year$^{-1}$). These values are from simulations covering the period from 2003 to 2010 of eight trees of the second simulation with initial DBH: 5cm, 15cm, 30cm and 40cm.

| DBH initial | NPP | | GPP | | LAI | | Rauto | |
|---|---|---|---|---|---|---|---|---|
| | Mean | sd | Mean | sd | Mean | sd | Mean | sd |
| 5 | 756.0 | 130.1 | 1523.6 | 161.4 | 5.3 | 0.6 | 767.5 | 140.6 |
| 15 | 756.5 | 124.8 | 1543.0 | 157.3 | 5.6 | 0.5 | 786.5 | 109.2 |
| 30 | 761.1 | 127.6 | 1569.7 | 156.6 | 6.1 | 0.6 | 808.6 | 96.5 |
| 40 | 742.0 | 134.0 | 1584.8 | 157.2 | 6.6 | 0.6 | 842.9 | 101.5 |
