## Supplementary material for "Comparing statistical and mechanistic models to identify the drivers of mortality within a rear-edge beech population": Suplementary appendices

### Appendix 1: CASTANEA model, calibration and simulation design

#### 1 - Modelling late frost damage

Here, we used the UniForc model a one-phase model, describing the cumulative effect of forcing temperatures on bud development during the ecodormancy phase (Chuine et al., 1999). Budburst occurs when the accumulated rate of forcing $R_f$ (Eqn. 1) reaches F[illegible]:

$$\sum_{d=t_0}^{t_f} R_f(T_d) \geq F$$

$R_f$ is calculated as a sigmoid function

$$\frac{1}{1 + e^{-d_t}(T_d - T_{50})}$$

with $d_T$ the positive slope and $T_{50}$ the mid-response temperature of the sigmoid function.

The level of frost damage was estimated using the following equations :

$$leafFrostDamage(t+1) = min\left(1; leafFrostDamage(t) + \frac{1}{1 + exp(BF * frostHardiness - T_{min})}\right)$$

With $T_{min}$ the minimal daily temperature, DL the day length and BF is a factor estimated as following:

$$BF = -0.3 - 1.5 \times exp(0.1 \times frostHardiness)$$

The frost hardiness was calculated as following:

$$frostHardiness(t+1) = frostHardiness(t) \times \frac{4}{5} + \frac{1}{5} \times \left(FH_{minfe} + CR \times (dFHti + dFHpi)\right)$$

$$With \begin{cases} if(T_{min} > T_{e1}) dFHti = 0 \\ if(T_{min} < T_{e1} \wedge T_{min} > T_{e2}) dFHti = if(T_{min} < T_{e2}) dFHti = FHtfemax \end{cases} FHtfemax - \frac{FHtfemax}{T_{e1} - T_{e2}} \times (T_{min} - T_{e2})$$

$$With \begin{cases} if(DL > NL2) dFHpi = 0 \\ if(DL < NL1) dFHpi = FHpfemax \\ if(DL < NL2 \wedge DL > NL1) dFHpi = FHpfemax - \frac{FHpfemax}{NL1 - NL2} \times (DL - NL2) \end{cases}$$

Then the effect of these damages was modeled according to three modalities: no effect, an effect with a definite decrease of the LAI, a decrease of the LAI followed by a new leaf flush. When unfolding leaves are affected by frost, decreased of leaf area index (LAI) is simulated every day using the following equation:

$$LAI = LAI_{initial}(1 - leafFrostDamage)$$

#### 2 - Calibration

We simulated a population of 100 trees representing the mean (and possibly the variability) in individual characteristics observed in La Massane in terms of height-diameter allometry (fixed), diameter (variable), leaf area index (fixed) and budburst phenology (variable).

The values of DBH were randomly drawn for each mean individual in a gamma law with mean and standard deviation measured in the whole population.

Budburst was simulated following equation 1-3 detailed in Oddou-Muratorio and Davi (2014), with the inter-individual variability in budburst date generated by varying a single parameter, FcritBB, the critical value of the state of forcing, which is most commonly referred to as the temperature sum required for budburst. We considered that the population was composed of 10 trees with a low FcritBB-value (early budburst), and of 90 trees with a the FcritBB-value observed for beech (“normal” budburst).

We also simulated a range of SWCa conditions representing the variability in SWCa in the site (TableA2.1). Height-diameter allometry (agF) was computed by simple linear regression (agF =1.26746 ). The LAI-value measured in La Massane is 4.48.

Inventory files for CASTANEA simulation are available in the Zenodo repository : https://doi.org/10.5281/zenodo.3519315 .

**Table A2.1 Values of parameters used for simulations at forest stand-level.** Distribution law used to draw parameters values (S.Law); respective mean and standard deviation (sd) for the Gaussian law; respective shape and rate of the Gamma law. Units of the variables.

| Variables | S.Law | Mean | sd | shape | rate | units |
|---|---|---|---|---|---|---|
| **SWCa** | Gaussian | 167.90 | 22.88 | - | - | |
| **Fcrit for trees with early budburst** | Gaussian | 41 | 5 | - | - | |
| **Fcrit for other trees** | Gaussian | 33 | 5 | - | - | |
| **DBH** | Gamma | - | - | 0.80 | 0.06 | cm |

#### 2- Validation

**Method:** To evaluate CASTANEA simulations, we used ring width profiles measured on 100 beech trees from the study site with contrasted size and defoliation characteristics. Cores were extracted in February 2016 at 1.30m above ground. After sanding, cores were scanned at 1200 dpi. Boundary rings were read using. Nine trees were excluded due to fungal contamination presence or unreadable ring width (indistinguishable RW or friable wood). Ring width were measured, and each individual series was cross-dated to check for missing rings and dating errors using the softwares Cdendro 9.0 and CooRecorder v 9.0 (Cybis Elektronik & Data AB. Sweden).

**Results and discussion:** Annual ring widths simulated by CASTANEA over the period from 1959 to 2015 were significantly correlated to the observed ring widths with an $R^2$ of 0.58 (p.value = 2.7e-6). (Fig. A2.1). Beyond this global correlation, two discrepancies between simulated and observed growth were noticeable (Fig. A2.2). CASTANEA simulated a trend in decreasing ring width, which was not mirrored in observed ring widths. Moreover, during two periods (from 1981 to 1992 and from 2000 to 2015), CASTANEA simulated depressions in ring widths, which were not mirrored in observed ring widths. This can be due to a slackening of competition in the observed population due for instance to tree mortality. Indeed, CASTANEA does not account for changes in tree dominance status, which can affect their current carbon balance and hence their growth.

**Figure A2.1 observed (x-axis) vs simulated ring width (y-axis) from 1960 to 2015.** Full line is the first bissectrice. Simulated ones are overestimated but the correlation of variations is significant 0.58 (pval= $2.66.10^{-6}$) .

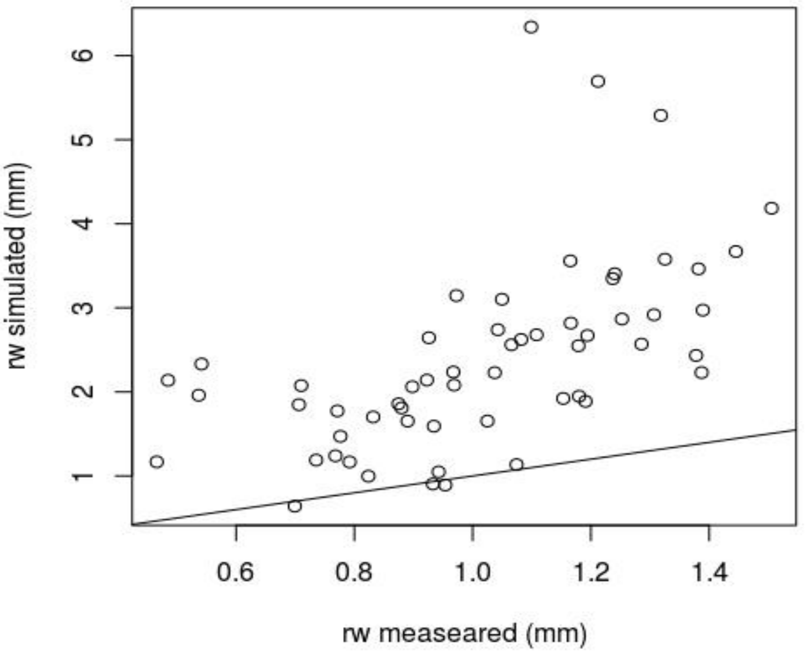

**Figure A2.2 observed Ring width (grey) vs simulated ring width (blue) from 1960 to 2015.** Grey dotted line is the mean observed ring width chronology. The grey envelope corresponds to the maximum and minimum ring width observed. Blue line is the mean simulated ring width. Blue envelope is the maximum and minimum simulated ring width.

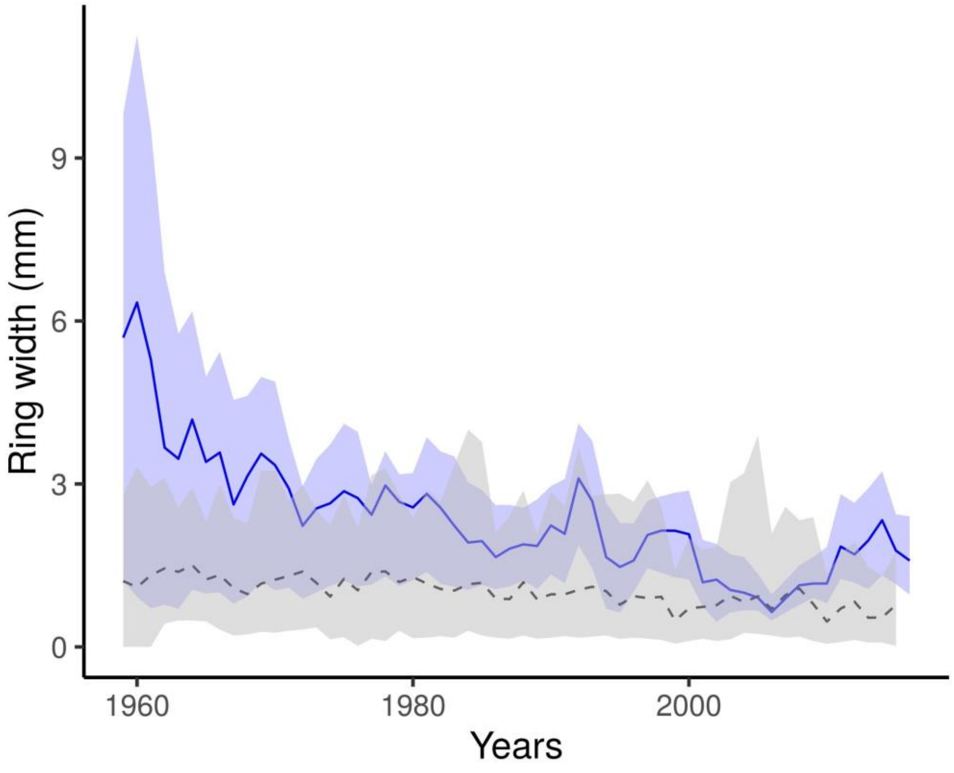

### Appendix 2: Beta-regression model for the temporal variations in the rate of mortality at forest stand level

This appendix details the results of a beta-regression model, which we also used to investigate the effects of climate on the inter-annual variations of mortality rate at forest stand-level. While in the main part of the manuscript we investigate the correlation between the response variables simulated by CASTANEA (NLF, PLC, BoR) and the mortality rate, here, we used as predictor variables only climatic compound variables computed from climatic series.

**Method:** Beta-regression models predict a response variable varying between [0; 1] and account for features like heteroscedasticity or asymmetry, which are commonly obtained in time-series of annual mortality rates. We investigated the following model for mortality at year *n* (with *n* varying from 2004 to 2016):

$$\tau_n = SPEI1\_dryVg_n \times SPEI3\_JJA_n \text{ (Equation A2.1)},$$

where SPEI1_dryVg is lowest monthly value of the Standardised Precipitation-Evapotranspiration Index (SPEI) during the vegetative season; and SPEI3_JJA is cumulated SPEI from June to August computed over three months. The SPEI is a multiscalar drought index based on climatic data varying between -2 and 2. It can be used for determining the onset, duration and magnitude of drought conditions with respect to normal conditions. We used the package SPEI (Beguería & Vicente-Serrano, 2017) to compute monthly SPEI-values from 1970 to 2015 considering two possible period of references for the “normal conditions” (respectively one and three month before the considered period). We gave the same weight to all previous months and used a uniform distribution to do it. Moreover, we modified the calibration of evapotranspiration (ETP), adapted to crop in SPEI package; we used instead the ETP estimation provided by the mechanistic model (CASTANEA). Then, for each of the two monthly SPEI series, we computed two SPEI variables for each year from 1970 to 2015. SPEI3_JJA is the mean SPEI computed over June, July and August, taking into account the precipitation of from April to August. SPEI1_dryVg is the minimum SPEI reach over April to August, taking into account the precipitation from April to August. As the number of observations (years) was low, we focussed on these two predictors expected to cause mortality based on beech ecology (Bréda et al. 2006).

Beta-regression was fitted with the R package ‘betareg’ (Cribari-Neto and Zeileis 2010). The variables were scaled before fitting the model. Model validity was checked based on the leverage points (i.e. points having a greater weight than expected by chance) with the Cook's distance (Cook distance < 0.5 indicate no leverage). We evaluated the goodness-of-fit with the Brier test score (Brier 1950). We evaluated the sensitivity and specificity of the model using the receiver operating characteristic (ROC) curve.

**Results and discussion:** The beta-regression model revealed a significant impact of climate variables on the observed mortality rate at forest stand-level (Table A2.1). SPEI1_dryVg, SPEI3_JJA and their interactions explained 32% of the variation in mortality rate between years. For low values of SPEI3_JJA (i.e. during dry summers), mortality increased with decreasing SPEI1_dryVg (i.e., increasing drought intensity). However, non-expected interaction effects were observed for high values of SPEI3_JJA (i.e. during wet summers), where mortality increased with increasing SPEI1_dryVg (i.e., decreasing drought intensity).

**Table A2.1: Parameter estimates and related statistics for beta-regression model described by equation A2.1.** β is the maximum likelihood estimate, with its estimated error (SE), z-value, and associated p-value ( Pr(>|z|). OR is the odds ratio. VIF is the generalized collinearity diagnostic value (variance inflation factor). The precision parameter estimate (ψ=435.9) was statistically significant (p-value=0.01).

| Variables | β | SE | z-value | p-value | OR | VIF |
|---|---|---|---|---|---|---|
| **SPEI1_dryVg** | -0.15 | 0.16 | -0.96 | 0.33 | $8.60.10^{-1}$ | 1.16 |
| **SPEI3_JJA** | 0.37 | 0.16 | 2.25 | 0.02 | 1.46 | 3.51 |
| **SPEI1_dryVg:SPEI3_JJA** | 0.31 | 0.16 | 1.89 | 0.06 | 1.36 | 3.28 |

The model had a good validity (VIF < 4) and goodness-of-fit (0.32 of $R^2$) (Table A2.1, Figure A2.1)) VIF were computed with the package "car" (Fox & Weisberg, 2011). With the diagnostic plot (Figure A1.3), we did not observe any trends in the residues, which means that there is no detectable temporal autocorrelation. We observed that 3 years have a higher impact than expected randomly on the model prediction 2004, 2006 and 2015 ( ie. cook distance > 0.5). For the first two leverage points, this is not surprising since these two years have high mortality rates.

This approach found a significant relationship between the observed mortality rate and SPEI variables computed from climatic series. A major role of SPEI on mortality has already been found by Archambeau et al. (2019) in beech, by Davi and Cailleret (2017) in silver fir, and by Carnicer et al., (2011) in 12 European tree species. The model suggested that mortality was triggered by summer droughts, including both the pulse effects of severe drought (through SPEI1_dryVg) but also long-term effects of repeated droughts (through SPEI3_JJA). Our results additionally suggest an interaction between long-term and pulse effects of drought on mortality i.e. the risk of mortality increased more than the sum of risks predicted by each factor separately. However, the biological interpretation of some of these interaction effects was not evident. This may be due to the low number of observations, with only 14 years in the mortality survey. This may also confirm that integrative measurements of the response to stresses, such as the CVI proposed in the main document, allow a better understanding of when and how mortality occurs than purely stress-related climatic variables.

**Figure A2.1: Diagnostic plot of the beta-regression.** From the top left to the bottom right: Residuals vs. indices of observation; Cook's distance plot; Generalized leverage vs predicted values; Residuals vs linear predictor.

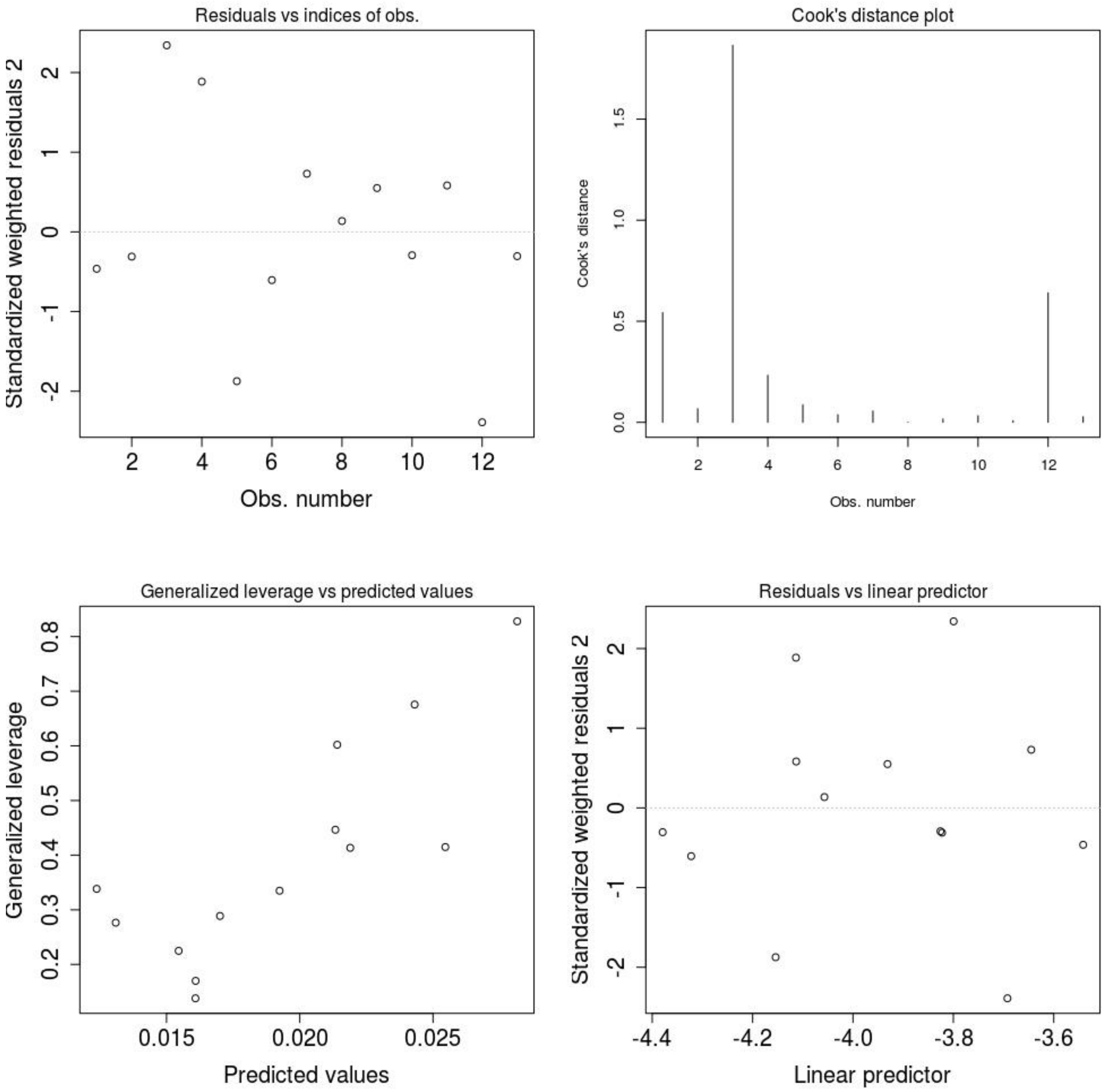

### Appendix 3: Logistic regression models for the probability of mortality at tree-level

This appendix aims to present the procedure of variable selection for the logistic regression model presented in equation 9 in the main document:

$$P_{mortality} = [DEFw + Fungi + Budburst + MBAI + (Nstem\ \textbf{OR}\ Compet_{intra}\ \textbf{OR}\ Compet_{intra+}\ \textbf{OR}\ Compet_{inter})] \times (DBH_{2002} + DBH_{2002}^2)$$

where $P_{mortality}$ is the individual probability of mortality; defoliation (DEFw), growth (MBAI), size ($DBH_{2002}$) and competition (Nstem and the Compet indices) factors were quantitative variables, while fungus presence (Fungi) and budburst phenology were categorical variables.

The section (1) presents the procedure for variable selection. Sections 2 to 4 present three alternative models where (2) DBH is considered as a categorical variable (instead of quantitative variable), (3) only non-clonal trees (ie with Nstem=1) are considered and (4) height in 2002 is considered instead of DBH.

#### 1- Variables and model selection

**Table A3.1 Comparison between models.** This table was built using MuMin package, and gives the name of the model; the variables present in the model, degree of freedom (df) ; the log-likelihood (logLik); the value of the information criterion used (AICc) ; the delta AIC (delta); and 'Akaike weight' (weight).

| Model Name | Variables | df | logLik | AICc | delta | weight |
|---|---|---|---|---|---|---|
| **"Nstem"** | [DEFw+Fungi+Budburst+MBAI+ Nstem ] ×( DBH2002+ DBH2002²) | 18 | -1630.55 | 3297.25 | 0 | 0.75 |
| **"intra+"** | [DEFw+Fungi+Budburst+MBAI+ Competintra+]×( DBH2002+ DBH2002²) | 18 | -1631.73 | 3299.62 | 2.39 | 0.23 |
| **"intra"** | [DEFw+Fungi+Budburst+MBAI+ Competintra]×( DBH2002+ DBH2002²) | 18 | -1634.07 | 3304.30 | 8.20 | 0.012 |
| **"tot"** | [DEFw+Fungi+Budburst+MBAI+ Compettot]×( DBH2002+ DBH2002²) | 18 | -1634.65 | 3305.46 | 9.30 | 0.007 |

**Table A3.2 Variables effects in the best model ("Nstem") and model validation. Df** are the degrees of freedom. **Dev** is the deviance associated. **AIC** is the AIC of the model if the variable is removed. **LRT** is the likelihood ratio test between the complete model and the model with the variable removed, with its associated p-value. **GVIF$^{1/2df}$** is the generalized collinearity diagnostic value taking into account the different dimensions of the variables with Df. The Brier score of the model is 0.11 and the goodness of fit (GoF) computed with the roc curve (area under the curve) is 0.82.

| Variables | Df | Dev. | AIC | LRT | p-value | GVIF | GVIF1/2df |
|---|---|---|---|---|---|---|---|
| **DEFw** | 1 | 4346.2 | 4380.2 | 1085.06 | < 2.20e-16 | 1.23 | 1.11 |
| **Fungi** | 1 | 3276.2 | 3310.2 | 15.15 | 9.91e-05 | 1.50 | 1.23 |
| **Budburst** | 1 | 3281.7 | 3315.7 | 20.64 | 5.55e-06 | 1.02 | 1.01 |
| **MBAI** | 1 | 3302.9 | 3336.9 | 41.80 | 1.01e-10 | 1.54 | 1.24 |
| **Nstem** | 1 | 3272.2 | 3306.2 | 11.08 | 8.73e-4 | 1.08 | 1.04 |
| **(DBH2002+DBH2002²)** | 2 | 3272.6 | 3304.6 | 11.46 | 3.25e-3 | 40.46 | 2.52 |
| **DEFw:(DBH2002+DBH2002²)** | 2 | 3273.9 | 3305.9 | 12.85 | 1.62e-3 | 6.64 | 1.61 |
| **Fungi:(DBH2002+DBH2002²)** | 2 | 3262.9 | 3294.9 | 1.79 | 0.41 | 5.74 | 1.55 |
| **Budburst:(DBH2002+DBH2002²)** | 2 | 3261.2 | 3293.2 | 0.07 | 0.97 | 1.34 | 1.08 |
| **MBAI:(DBH2002+DBH2002²)** | 2 | 3265.3 | 3297.3 | 4.26 | 0.12 | 2.22 | 1.22 |
| **Nstem:(DBH2002+DBH2002²)** | 2 | 3261.3 | 3293.3 | 0.21 | 0.90 | 14.95 | 1.97 |

**Fig A3.1: ROC curve.** Percent of true positives (sensitivity 49%) against percent of false positives (1 – specificity 51% ). The percentage of true negatives is 96% and the percentage of false negatives is 4%.

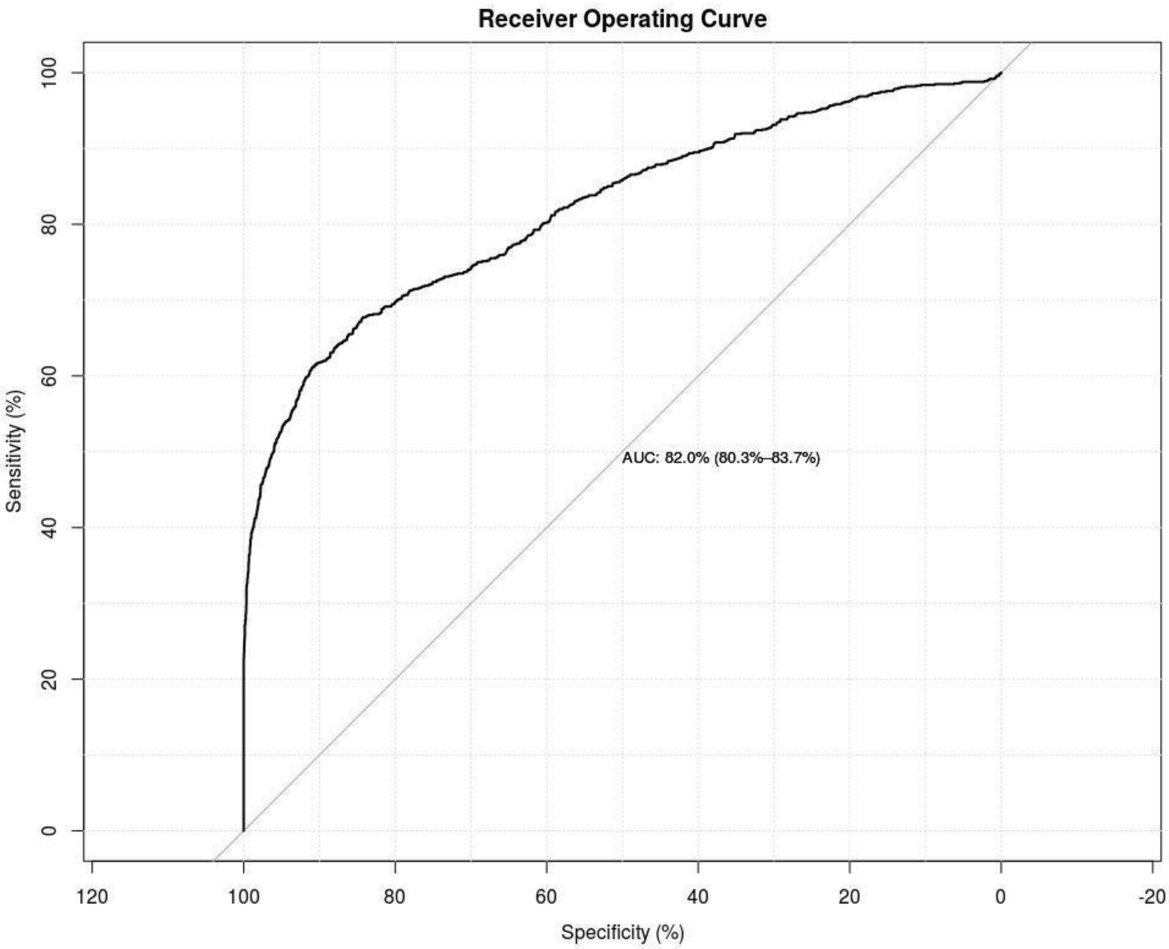

**Figure A3.2: Cook's distance plot.** Stand Pearson residuals and their leverage. If the point is out of 0.5, it has a big influence in the model.

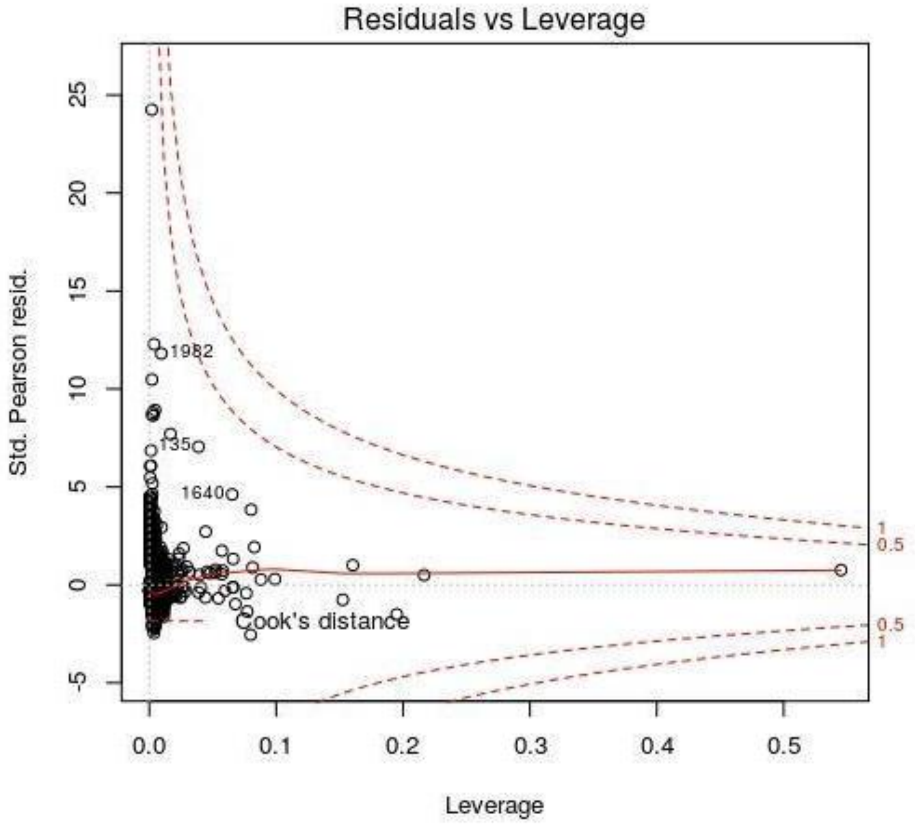

#### 2- Alternative model where DBH is considered as a class qualitative variable

To take into account the U-shaped relationship between mortality and DBH, we also tested a model with DBH considered as a 4-level categorical variable. We distinguished four DBH-classes: Class 10-20cm with 1280 individuals, class 21-30cm with 1312 individuals, class 31-40cm with 411 individuals and class >40cm with 200 individuals. This model wrote as follows :

$P_{mortality}$ = [DEFcum + Fungi **+** Budburst**+** MBAI **+**
(Nstem **OR** $Compet_{intra}$**OR** $Compet_{tot}$ **OR** $Compet_{intra+}$)] * Class_DBH (equation A3.2)

We applied the same procedure as in the main article to select first the best competition variable, and then use the step AIC procedure to select the best model.

**Table A3.3: Parameter estimates and related statistics in the best model (eq. A3.2).** β is the maximum likelihood estimates and SE its estimated error. We give the z value and the associated p-value of the test that β is significantly different from zero. OR is the estimated odds ratios.

| Variables | β | SE | z value | p-value | OR |
|---|---|---|---|---|---|
| **DEFw** | 7.512 | 0.364 | 20.653 | < 0.001 | 1830.1 |
| **class_DBH21-30** | -0.253 | 0.172 | -1.471 | 0.141173 | 0.776 |
| **class_DBH31-40** | -1.102 | 0.302 | -3.646 | < 0.001 | 0.332 |
| **class_DBH>40** | -0.204 | 0.467 | -0.436 | 0.663 | 0.816 |
| **MBAI** | -0.077 | 0.017 | -4.606 | < 0.001 | 0.926 |
| **Budburst (Early)** | 0.783 | 0.172 | 4.544 | < 0.001 | 2.189 |
| **Nstem** | 0.122 | 0.036 | 3.388 | < 0.001 | 1.130 |
| **Fungi (Presence)** | 0.493 | 0.143 | 3.443 | < 0.001 | 1.637 |
| **class_DBH21-30:MBAI** | -0.037 | 0.027 | -1.373 | 0.169 | 0.964 |
| **class_DBH31-40:MBAI** | 0.081 | 0.020 | 4.142 | < 0.001 | 1.085 |
| **class_DBH>40:MBAI** | -0.059 | 0.038 | -1.534 | 0.125 | 0.943 |
| **DEFw:class_DBH21-30** | -1.575 | 0.560 | -2.813 | < 0.001 | 0.207 |
| **DEFw:class_DBH31-40** | -2.100 | 0.822 | -2.556 | < 0.001 | 0.122 |
| **DEFw:class_DBH>40** | 0.243 | 0.025 | 3.4372 | 0.254 | -1.416 |

**Table A3.4 Variables effects in the best model (eq. A3.2) and model validation.** Df are the degree of freedom. Dev. is the deviance associated. AIC is the AIC of the model if the variable is removed. LRT is the likelihood ratio test between the complete model and the model with the variable removed, with its associated p-value. GVIF1/2df is the generalized collinearity diagnostic value taking into account the different dimensions of the variables with Df. The Brier score of the model is 0.11 and the goodness of fit computed with the roc curve (area under the curve) is 0.83.

| Variables | Df | Dev. | AIC | LRT | p-value | GVIF1/2df |
|---|---|---|---|---|---|---|
| **DEFw** | 1 | 3975.9 | 4003.9 | 725.57 | < 0.001 | 1.523 |
| **class_DBH** | 3 | 3266.7 | 3290.7 | 16.39 | < 0.001 | 1.798 |
| **MBAI** | 1 | 3279.7 | 3307.7 | 29.42 | < 0.001 | 2.258 |
| **Budburst** | 1 | 3269.6 | 3297.6 | 19.31 | < 0.001 | < 0.001 |
| **Nstem** | 1 | 3261.2 | 3289.2 | 10.88 | < 0.001 | 1.005 |
| **Fungi** | 1 | 3261.9 | 3289.9 | 11.57 | < 0.001 | 1.084 |
| **class_DBH:MBAI** | 3 | 3291.1 | 3315.1 | 40.75 | < 0.001 | 1.595 |
| **DEFw:class_DBH** | 3 | 3261.7 | 3285.7 | 11.34 | 0.0100 | 1.848 |

#### 3- An alternative model with competition index as a proxy for competition.

To test the robustness of the model to the choice of the competition predictor, we first reduced the data set by excluding all trees belonging to a coppice (keeping only tree with Nstem =1). Then, we fitted the following model

$$P_{mortality} = [\text{DEFcum} + \text{Fungi} + \text{Budburst} + \text{MBAI} + (\text{Compet}_{intra}\ \textbf{OR}\ \text{Compet}_{tot})] * (\text{DBH} + \text{DBH}^2)$$

(equation A3.3)

We applied the same step AIC procedure as in the main article to select the best model The fit of this model revealed that all the variables had a significant effect on the probability of mortality except competition. Moreover, these effects were consistent with those estimated in the main part of this manuscript.

**Table A3.5: Parameter estimates and related statistics for the best model (eq. A3.3).** β is the maximum likelihood estimates and SE its estimated error. We give the z value and the associated p-value of the test that β is significantly different from zero. OR is the estimated odds ratios.

| Variables | β | SE | z value | p-value | OR |
|---|---|---|---|---|---|
| **DEFw** | 6.912 | 0.263 | 26.312 | < 0.001 | 857.030 |
| **DBH2002** | -12.189 | 5.141 | -2.371 | < 0.001 | 0.927 |
| **DBH2002²** | 22.306 | 4.949 | 4.507 | 0.104 | 1.6 10-4 |
| **MBAI** | -0.060 | 0.011 | -5.573 | < 0.001 | 5.76 106 |
| **Budburst** | 0.842 | 0.172 | 4.899 | < 0.001 | 2.413 |
| **Fungi** | 0.580 | 0.142 | 4.092 | < 0.001 | 1.919 |
| **DEFw: DBH2002** | -56.404 | 14.792 | -3.813 | 0.03 | 3.9 10-16 |
| **DEFw: DBH2002²** | 32.591 | 16.851 | 1.934 | 0.02 | 537.8 |
| **DBH2002:MBAI** | 0.381 | 0.454 | 0.839 | 0.551 | 0.687 |
| **DBH2002²:MBAI** | -0.835 | 0.443 | -1.884 | 0.105 | 0.379 |

**Table A3.6 Variables influences in the best model (eq. A3.3) and model validation.** Df is the number of degree of freedom. Dev. is the deviance associated. AIC is the AIC of the model when the variable is removed. LRT is the likelihood ratio test between the complete model and the model with the variable removed, with its associated p-value. GVIF1/2df is the generalized collinearity diagnostic value taking into account the different dimensions of the variables with Df. The Brier score of the model is 0.24 and the goodness of fit computed with the roc curve (area under the curve) is 0.82.

| Variables | Df | Dev. | AIC | LRT | Pr(>Chi) | GVIF | GVIF1/2df |
|---|---|---|---|---|---|---|---|
| **DEFw** | 1 | 2915.6 | 2935.6 | 713.39 | < 0.001 | 1.258 | 1.122 |
| **MBAI** | 1 | 2236.2 | 2256.2 | 34.01 | < 0.001 | 1.510 | 1.229 |
| **DBH2002** | 2 | 2215.9 | 2233.9 | 13.71 | 0.001 | 5.267 | 1.515 |
| **Budburst-early** | 1 | 2217.3 | 2237.3 | 15.14 | < 0.001 | 1.019 | 1.010 |
| **Fungi-T** | 1 | 2215.4 | 2235.4 | 13.24 | < 0.001 | 1.223 | 1.106 |
| **DEFw: DBH2002²** | 2 | 2213.9 | 2231.9 | 11.68 | 0.003 | 13.795 | 1.927 |
| **MBAI: DBH2002** | 2 | 2207 | 2225 | 4.78 | 0.0916 . | 10.346 | 1.793 |

**4 - Alternative model with Height in 2002 as a proxy of size**

To test he robustness of the model to the choice of the size predictor, we also a considered a model where size was measured by height instead of DBH:

Pmortality = [DEFw + Fungi + Budburst+ MBAI +

(Nstem OR $Compet_{intra}$ OR $Compet_{tot}$ OR $Compet_{intra+}$)] × $H_{2002}$ (equation A3.4)

The height in 2002 ($H_{2002}$) varied between 1.5m and 26m and was measured only for a subset of 1199 trees. We applied the same procedure as in the main article for the correlated competition variables. Then, we used the step AIC procedure to select the best model.

The fit of this model revealed less variables with a significant effect than previously. This is probably due to the lower number of individual observations and also to the fact that the measured trees are almost all dominant. The significant effects were consistent with those estimated in the main part of this manuscript.

**Table A3.7: Parameter estimates and related statistics for the best model (eq. 3.4).** β is the maximum likelihood estimates and SE its estimated error. We give the z value and the associated p-value of the test that β is significantly different from zero. OR is the estimated odds ratios.

| Variables | β | SE | z value | p-value | OR |
|---|---|---|---|---|---|
| **DEFw** | 6.657 | 0.481 | 13.831 | < 0.001 | 778.50 |
| **H_2002** | -0.215 | 0.038 | -5.642 | < 0.001 | 0.81 |
| **MBAI** | -0.059 | 0.018 | -3.237 | 0.0012 | 0.94 |
| **Budburst (Early)** | 0.791 | 0.308 | 2.568 | 0.0102 | 2.21 |

**Table A3.8 Variables influences in the best model (eq. A3.4) and model validation.** Df is the number of degree of freedom. Dev. is the deviance associated. AIC is the AIC of the model when the variable is removed. LRT is the likelihood ratio test between the complete model and the model with the variable removed, with its associated p-value. GVIF1/2df is the generalized collinearity diagnostic value taking into account the different dimensions of the variables with Df. The Brier score of the model is 0.10 and the goodness of fit computed with the roc curve (area under the curve) is 0.85.

| Variables | Df | Dev. | AIC | LRT | p-value | GVIF |
|---|---|---|---|---|---|---|
| **DEFw** | 1 | 1105.28 | 1113.28 | 295.34 | < 0.001 | 1.05 |
| **H2002** | 1 | 845.77 | 853.77 | 35.823 | < 0.001 | 1.04 |
| **MBAI** | 1 | 823.44 | 831.44 | 13.494 | < 0.001 | 1.06 |
| **Budburst** | 1 | 816.08 | 824.08 | 6.129 | 0.013 | 1.01 |

### Appendix 4: Survival analysis of the probability of mortality at tree- and year-levels

This appendix aims to present an attempt to fit a survival analysis model to our data set. Survival analysis typically consists in analysing the relationships between the time that elapsed before mortality occurred on the one hand, and the measured covariates that may be associated with that quantity of time on the other hand. This is not exactly similar to measuring the effect of each covariate on the risk of mortality, as presented in the main part of this manuscript.

#### Method

We created a new data table where each individual is repeated (several lines) as many years as those during which it is observed as alive. For the fungi variable (FungiEvent), once visible the status changes from 0 to 1. Two types of defoliation variables were considered: Def_ev is the defoliation mark measured for a given year (0= non defoliated; 1=defoliated). DEF_cum is the sum of defoliation marks up to a given year (DEF_cum varies between 0 and 9). The model writes as follows:

Pmortality = [DEF_cum + Def_ev + Fungi + Budburst+ MBAI +
(Nstem OR $Compet_{intra}$ OR $Compet_{tot}$ OR $Compet_{intra+}$)] * ($DBH+DBH^2$) (equation 4)

We applied the same procedure as in the main article to select the best competition variable. Then, we used the step AIC procedure to select the best model.

#### Results

The fit of the survival model revealed the same effects as found with the logistic regression model presented in the main text (Table A3.9 and Table 2). In particular, the relative probability of mortality increased with increasing competition (Nstem) and decreased with increasing growth; it was also higher for trees with an earlier budburst as compared to others, and for trees bearing fungi fructifications than for others. Interestingly, we observed different effects of the two variables related to defoliation: On the one hand, increasing cumulated defoliation (DEF_cum) increased the risk of mortality. On the other hand, increasing single-year defoliation (Def_ev) decreased the risk of mortality. Almost all years had a positive or negative effect compared to 2003 (except 2010 or the lack of data, did not allow an assessment of the effects).

However, this survival analyses model did not seem fully reliable, because of a major departure from classical assumptions of survival analyses. Indeed, survival analysis following the classical cox model assume proportional hazards, ie the unique effect of a defoliation mark equal to 1 instead of 0 would for instance double the probability of mortality at every year t. However, the effect of defoliation on mortality is likely to vary among years depending on climate, and this cannot easily be taken into account in survival analyses. None of the variables respected the proportional-hazards (PH) assumption (table A3.10).

**Table A3.9: Parameter estimates and related statistics for the survival analysis model.** β is the maximum likelihood estimates and SE its estimated error. OR the estimated odds ratios. We give the z value and the associated p-value of the test that β is significantly different from zero.

| Variables | β | SE | OR | z | p-value |
|---|---|---|---|---|---|
| **Fungi_event** | $5.057\ 10^{-1}$ | $9.558\ 10^{-2}$ | 1.658 | 5.291 | < 0.001 |
| **Def_cum** | $6.945\ 10^{-1}$ | $2.406\ 10^{-2}$ | 2.003 | 28.869 | < 0.001 |
| **DBH2002** | -8.615 | $1.093\ 10^{-1}$ | $1.813\ 10^{-4}$ | -0.788 | 0.431 |
| **DBH2002²** | $4.772\ 10^{1}$ | 8.965 | $5.3\ 10^{20}$ | 5.323 | < 0.001 |
| **Budburst** | $5.972\ 10^{-1}$ | $1.214\ 10^{-1}$ | 1.817 | 4.921 | < 0.001 |
| **MBAI** | $-9.233\ 10^{-2}$ | $8.846\ 10^{-3}$ | $9.118\ 10^{-1}$ | -10.438 | < 0.001 |
| **Nstem** | $9.858\ 10^{-2}$ | $2.313\ 10^{-2}$ | 1.104 | 4.261 | < 0.001 |
| **Def_ev** | -1.369 | $1.200\ 10^{-1}$ | $2.543\ 10^{-1}$ | -11.412 | < 0.001 |
| **Year 2004** | $4.702\ 10^{-1}$ | $1.396\ 10^{-1}$ | 1.600 | 3.368 | < 0.001 |
| **Year 2005** | $6.228\ 10^{-2}$ | $1.464\ 10^{-1}$ | 1.064 | 0.425 | 0.671 |
| **Year 2006** | $2.633\ 10^{-1}$ | $1.378\ 10^{-1}$ | 1.301 | 1.911 | 0.056 |
| **Year 2007** | $-1.533\ 10^{-1}$ | $1.434\ 10^{-1}$ | $8.58\ 10\text{-}^{1}$ | -1.069 | 0.285 |
| **Year 2008** | -1.844 | $2.160\ 10^{-1}$ | $1.581\ 10^{-1}$ | -8.541 | < 0.001 |
| **Year 2009** | -1.660 | $1.984\ 10^{-1}$ | $1.901\ 10^{-1}$ | -8.370 | < 0.001 |
| **Year 2010** | NA | 0.000 | NA | NA | NA |
| **Year 2011** | -1.019 | $1.691\ 10^{-1}$ | $3.609\ 10^{-1}$ | -6.025 | < 0.001 |
| **Year 2012** | $-7.465\ 10^{-1}$ | $1.566\ 10^{-1}$ | $4.740\ 10^{-1}$ | -4.766 | < 0.001 |
| **Year 2013** | $-9.312\ 10^{-1}$ | $1.652\ 10^{-1}$ | $3.941\ 10^{-1}$ | -5.636 | < 0.001 |
| **Year 2014** | -1.029 | $1.705\ 10^{-1}$ | $3.574\ 10^{-1}$ | -6.037 | < 0.001 |
| **Year 2015** | -1.361 | $1.862\ 10^{-1}$ | $2.564\ 10^{-1}$ | -7.308 | < 0.001 |
| **Year 2016** | -1.923 | $2.069\ 10^{-1}$ | $1.462\ 10^{-1}$ | -9.291 | < 0.001 |
| **Defcum:DBH2002** | $-1.793\ 10^{+1}$ | 4.487 | $1.628\ 10^{-8}$ | -3.996 | < 0.001 |
| **Def_cum: DBH2002²** | 7.488 | 3.687 | $1.787\ 10^{3}$ | 2.031 | 0.042 |
| **DBH2002:Def_ev** | $-9.782\ 10^{+1}$ | $3.18\ 10^{1}$ | $3.30\ 10^{-43}$ | -3.069 | 0.0021 |
| **DBH2002²:Def_ev** | $3.708\ 10^{+1}$ | $1.975\ 10^{1}$ | $1.2\ 10^{16}$ | 1.877 | 0.0604. |

**Table A3.10: Table of the result of the proportional-hazards (PH) assumption test.** Rho is the correlation coefficient between transformed survival time and the scaled Schoenfeld residuals, chisq the chi-square, and the two-sided p-value.

| Variables | rho | chisq | p-value |
|---|---|---|---|
| **Fungi (Presence)** | 0.06380 | 4.81 | $2.82\ 10^{-2}$ |
| **Def_cum** | -0.23724 | $5.25\ 10^{1}$ | $4.32\ 10^{-13}$ |
| **DBH2002** | -0.05881 | 4.25 | $3.94\ 10^{-2}$ |
| **DBH2002²** | 0.03750 | 1.41 | $2.35\ 10^{-1}$ |
| **Budburst (Early)** | 0.04496 | 2.16 | $1.42\ 10^{-1}$ |
| **MBAI** | 0.00513 | $1.54\ 10^{-1}$ | $6.94\ 10^{-1}$ |
| **Nstem** | 0.00117 | 1.15 10-03 | $9.73\ 10^{-1}$ |
| **Def_ev** | 0.13848 | $2.06\ 10^{1}$ | $5.63\ 10^{-6}$ |
| **Year 2004** | 0.04662 | 2.34 | $1.26\ 10^{-1}$ |
| **Year 2005** | 0.14269 | $2.17\ 10^{1}$ | $3.22\ 10^{-6}$ |
| **Year 2006** | 0.24548 | $5.94\ 10^{1}$ | $1.29\ 10^{-14}$ |
| **Year 2007** | 0.36127 | $1.29\ 10^{2}$ | < 0.001 |
| **Year 2008** | 0.31053 | $9.86\ 10^{1}$ | < 0.001 |
| **Year 2009** | 0.35782 | $1.32\ 10^{2}$ | < 0.001 |
| **Year 2010** | NA | NA | NA |
| **Year 2011** | 0.43059 | $1.89\ 10^{2}$ | < 0.001 |
| **Year 2012** | 0.49930 | $2.76\ 10^{2}$ | < 0.001 |
| **Year 2013** | 0.53221 | $3.04\ 10^{2}$ | < 0.001 |
| **Year 2014** | 0.56709 | $3.43\ 10^{2}$ | < 0.001 |
| **Year 2015** | 0.56088 | $3.29\ 10^{2}$ | < 0.001 |
| **Year 2016** | 0.53854 | $3.19\ 10^{2}$ | < 0.001 |
| **Def_cum:DBH2002** | 0.10397 | 8.84 | $2.94\ 10^{-3}$ |
| **Def_cum: DBH2002²** | -0.04641 | 1.81 | $1.79\ 10^{-1}$ |
| **DBH2002:Def_ev** | 0.02150 | $6.01\ 10^{-1}$ | $4.38\ 10^{-1}$ |
| **DBH2002²:Def_ev** | 0.00523 | $2.21\ 10^{-2}$ | $8.82\ 10^{-1}$ |
| **GLOBAL** | NA | $1.06\ 10^{3}$ | < 0.001 |
